# Sensor-mediated fine-tuning of siRNA levels is required for spermatogenic piRNA pathway function

**DOI:** 10.64898/2026.03.20.713214

**Authors:** Ha Meem, Alicia K. Rogers

**Affiliations:** Department of Biology, University of Texas at Arlington, Arlington, TX, 76019, USA

## Abstract

Small RNA pathways provide a robust and dynamic regulatory network that enables spatiotemporal regulation of the germline genome in response to environmental cues. The flexibility of RNA interference (RNAi)-mediated gene regulation and network architecture of these pathways requires molecular mechanisms that can fine-tune their regulatory potential and function to ensure proper execution of physiological processes, such as fertility. In *C. elegans*, we previously discovered a set of small RNA sensors that modulate the production of one class of small RNAs to adjust amplification resources based on cellular needs. These sensors maintain homeostatic levels of 22G-RNAs for the distinct RNAi branches that compete for resources in the *mutator* complex. Here we show this molecular feedback is essential for restricting expression of spermatogenic transcripts to an appropriate threshold during development and preventing spermiogenesis defects. Furthermore, we demonstrate 22G-RNA homeostasis is critical for proper meiotic progression in the germline and piRNA pathway function within pachytene germ cells. Together, our work reveals that RNAi homeostasis is critical for developmental and physiological processes, such as sperm-based fertility. Further, our findings show that small RNA pathway function is more than the sum of its parts and disrupting the ability to maintain homeostasis within the regulatory pathway itself leads to deleterious physiological consequences.

## INTRODUCTION

Maintenance of cellular homeostasis is critical to preserving cellular identity, and in germ cells, reproductive potential. Small RNA-mediated gene regulation, also termed RNA interference (RNAi), is a conserved mechanism that provides cellular homeostasis, silences foreign genetic elements such as transposons, and ensures appropriate gene expression in all animals. It is evident that RNAi pathways play a key role in dynamically regulating germline genes across eukaryotes^1-6^. However, to date, the mechanisms that maintain small RNA homeostasis and its role in physiological processes have been understudied. This is due to the difficulty in specifically perturbing homeostatic mechanisms without functionally disrupting core RNAi pathway components.

RNAi pathways rely on the RNA-induced silencing complexes (RISC) which regulate endogenous and exogenous transcript levels. Each RISC is made up of an Argonaute (AGO) protein and a loaded small RNA, typically 20∼30 nucleotide in length^7-10^. The small RNA provides sequence specificity for target transcripts while the AGO facilitates regulation of the target through transcriptional or post-transcriptional mechanisms^11^. Thus, we can define RNAi branches based on the AGO component of the RISC. The three different types of eukaryotic small RNAs, microRNAs (miRNAs), small interfering RNAs (siRNAs), and Piwi-interacting RNAs (piRNAs), differ based on their mode of biogenesis, AGO cofactor, and target transcripts^12^. Several mechanisms exist to generate small RNAs. For example, in *Caenorhabditis elegans*, the RNAse-III-like enzyme, Dicer, cleaves exogenous and endogenous long double-stranded RNAs (dsRNAs) to generate siRNAs referred to as “primary” siRNAs that are 26 nucleotides (nt) long with a 5’ guanine bias (26G-RNAs)^13,14^. In contrast, RNA polymerase II (Pol II) transcribes piRNA precursor transcripts that are capped and processed into mature piRNAs that are 21 nt long with a 5’ uridine bias (21U-RNAs)^15^. The 26G-RNAs and 21U-RNAs are loaded into distinct AGOs to form “primary RISC” that recognize target transcripts complementary to the loaded 26G or 21U RNA. Then, amplification of “secondary” siRNAs that are 22 nt long and have a 5’ guanine bias (22G-RNAs) is initiated by RNA-dependent RNA polymerases (RdRPs)^5,7,9,13,14,16^. This amplification step is essential for robust, heritable maintenance of RNAi-mediated regulation of the target^16-21^.

In *C. elegans*, amplification of several distinct RNAi pathway branches, such as ERGO-1, PRG-1 (the *C. elegans* PIWI-clade AGO), and RDE-1, rely on the *mutator* complex inside a perinuclear germ granule called the *Mutator* focus^16,22^. Thus, these RNAi branches compete for *mutator* complex resources. In contrast, other RNAi branches, such as the ALG-3/4 and CSR-1 branches, use *mutator*-independent 22G-RNA amplification mechanisms^16,22-25^. This network topology dictates that distinct RNAi branches sharing core molecular machinery in the *Mutator* focus are interdependent – disturbances to one branch can affect others. For example, reduction of one RNAi branch’s 26G-RNAs inputted into the *Mutator* focus would free-up amplification machinery resources leading to increased output of other RNAi branches’ 22G-RNAs. Supporting this model, we previously showed that depletion of upstream ERGO-1-class 26G-RNAs feeding into the *Mutator* focus triggered increased production of 22G-RNAs from the competing PRG-1 and RDE-1 RNAi branches^26^.

Proper coordination of RNAi-mediated gene regulation therefore requires delicate balancing to achieve homeostatic small RNA levels and AGO-loading fidelity. However, the mechanisms that maintain this balancing act remain largely unknown. Our previous work uncovered a feedback mechanism that maintains homeostatic levels of *mutator-*dependent 22G-RNAs. In this feedback mechanism, expression of *eri-6/7*, a helicase required for the production of ERGO-1-class 26G-RNAs, is regulated in a small RNA-dependent manner^26^ (**Figure 1A**). Functional ERI-6/7 protein is the translational product of a trans-spliced mRNA containing the *eri-6[a-d]* isoforms and *eri-7*^27^. The feedback mechanism regulates *eri-6/7* transcript levels through two independent sensor regions, *sensor of siRNAs-1* (*sosi-1*) and *eri-6[e-f],* embedded within an intron of *eri-6*. *Sosi-1* and *eri-6[e-f]* are targets of *mutator*-dependent 22G-RNAs. Low levels of 22G-RNAs complementary to the sensors trigger increased production of *eri-6[e-f]* isoforms, which cannot be trans-spliced to *eri-7* transcripts. This results in reduced mature, trans-spliced *eri-6/7* mRNAs, ERI-6/7 protein levels, and consequently, reduced biogenesis of ERGO-class 26G-RNAs. This cascade of events allows for increased production of non-ERGO-1-class 22G-RNAs by the *mutator* complex until sufficient levels of 22G-RNAs complementary to the sensors are reestablished.

**Figure 1.**
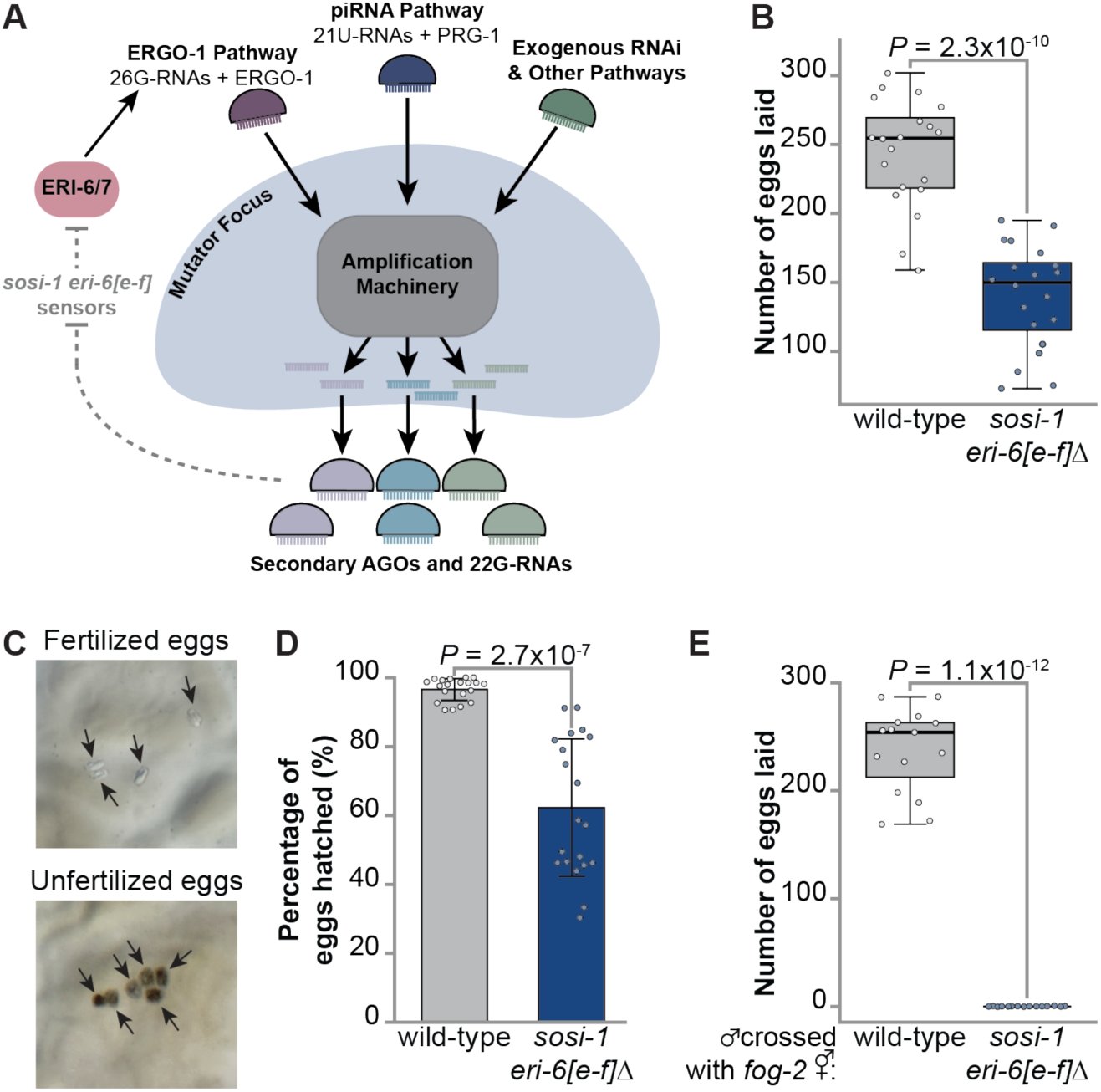
Sensor-mediated maintenance of homeostatic 22G-RNA levels is required for sperm-based fertility. **(A)** Schema of the competition of resources during *mutator-de*pendent amplification of different classes of 22G-RNAs and the role of *sosi-1* and *eri-6[e-f]* sensors in maintaining 22G-RNA homeostasis. Gray text and gray dashed lines indicate interactions that are ablated in *sosi-1 eri-6[e-f]Δ* mutants. **(B)** Total number of eggs laid by wild-type animals (gray) and *sosi-1 eri-6[e-f]Δ* mutants (blue). Circles represent number of eggs counted for individual hermaphrodites. Bolded midline indicates median value, box indicates the first and third quartiles, and whiskers represented the most extreme data points within 1.5 times the interquartile range, excluding outliers. **N** = 20. **(C)** Representative images of fertilized and unfertilized eggs. Black arrows point to eggs. **(D)** Total percentage of eggs hatched in wild-type animals (gray) and *sosi-1 eri-6[e-f]Δ* mutants (blue). Circles represent percentage of hatched eggs for individual hermaphrodites. Error bars indicate standard deviation. **N** = 20. (E) Box plots depicting total number of eggs laid for *fog-2(oz40)* hermaphrodites mated with wild-type males (gray) and *sosi-1 eri-6[e-f]Δ* males (blue). Circles represent hatched larvae from individual *fog-2* hermaphrodites after mating. Bolded midline indicates median value, box indicates the first and third quartiles, and whiskers represented the most extreme data points within 1.5 times the interquartile range, excluding outliers. **N** = 15. For **(B, D, E)** statistical significance was calculated by two-tailed Welch’s *t* test, mutants were compared to wild-type.

Our working model posits that changes in RNAi-mediated silencing of *sosi-1* and *eri-6[e-f]* fine-tunes ERGO-1 class 26G-RNA levels to balance the different RNAi branches that compete for the *mutator* complex. Due to the intronic location of the sensors, we were able to generate a mutant strain with the *sosi-1* and *eri-6[e-f]* sensor regions deleted (*sosi-1 eri-6[e-f]Δ*). Previously, we showed removal of the sensors does not disrupt *eri-6/7* function^26^. Thus, the *sosi-1 eri-6[e-f]1′* mutant is the first of its kind in any species that allows us to explore the role of RNAi homeostasis without altering the functionality of any RNAi pathway proteins. Here, we leverage the *sosi-1 eri-6[e-f]1′* mutant to explore the role of RNAi homeostasis in sperm-based fertility.

## RESULTS

### Sensor-mediated regulation of ERI-6/7 is required for sperm-based fertility

Small RNA pathways are critical for fertility across species, and many small RNA pathway mutants in *C. elegans* exhibit fertility defects^7,28,29^. To determine if sensor-mediated maintenance of 22G-RNA levels plays a role in fertility, we performed a brood size assay using wild-type (N2) and *sosi-1 eri-6[e-f]Δ* hermaphrodites. Synchronized L1 worms were plated on NGM plates and cultured at 20°C and individual L4 hermaphrodites were isolated prior to their brood size being assessed. We found *sosi-1 eri-6[e-f]Δ* mutants had significantly reduced brood sizes, laying roughly ∼42.52% fewer eggs compared to wild-type animals (**Figure 1B**). Further, we noticed that *sosi-1 eri-6[e-f]Δ* hermaphrodites lay more unfertilized eggs, which are noticeably darker than fertilized eggs, and had a significantly increased frequency of eggs that fail to hatch (35.95%) compared to wild-type animals (**Figures 1C and 1D**). Together, these results suggest *sosi-1 eri-6[e-f]Δ* hermaphrodites have a sperm-based fertility defect. To determine if removal of the sensors impacts sperm-based fertility, we mated *fog-2(oz40)* hermaphrodites, which are unable to produce their own sperm^30^, with wild-type and *sosi-1 eri-6[e-f]Δ* mutant males. Strikingly, *sosi-1 eri-6[e-f]*Δ males were unable to sire progeny when mated with *fog-2* hermaphrodites, confirming that the small RNA sensors are required for sperm-based fertility (**Figure 1E**). The reduced brood size in *sosi-1 eri-6[e-f]Δ* mutant hermaphrodites compared to the complete lack of progeny after mating with *sosi-1 eri-6[e-f]Δ* mutant males also suggests that male-derived sperm are more sensitive to perturbations in 22G-RNA homeostasis than hermaphrodite-derived sperm. This likely reflects differences in sperm development in hermaphroditic and male *C. elegans.* Taken together, our data indicate that the ability to maintain homeostatic 22G-RNA levels through the *sosi-1 eri-6[e-f]* sensors is critical for sperm-based fertility.

### Small RNA-mediated dysregulation of germline genes in the absence of *sosi-1* and *eri-6[e-f]* sensors

To understand how removal of the sensor-mediated feedback impacts fertility, we sought to identify genes with expression changes that correlated with the reduced sperm-based fertility of *sosi-1 eri-6[e-f]*Δ animals. We focused on the L4 developmental stage, when spermatogenesis occurs in the hermaphroditic germline of *C. elegans*. To this end, we generated mRNA-seq and size-selected small RNA-seq libraries from total RNA extracted from synchronized wild-type and *sosi-1 eri-6[e-f]*Δ L4 hermaphrodites (**Table S1**). Differential expression analyses of our mRNA-seq libraries found 938 genes were significantly up-regulated and 428 genes were significantly down-regulated in *sosi-1 eri-6[e-f]*Δ L4s compared to wild-type L4 animals. We then assessed changes in germline-specific (sex-neutral, oogenic, and spermatogenic) genes. We found sex-neutral germline genes and oogenic genes were significantly down-regulated at the mRNA level (**Figure 2A).** Indeed, sex-neutral germline genes and oogenic genes were enriched amongst the list of down-regulated genes and depleted from the list of up-regulated genes (**Figures S1A and S1B**). In contrast, spermatogenic genes were significantly up-regulated and enriched amongst the list of up-regulated genes and depleted from the list of down-regulated genes (**Figures S1A, S1B, and 2A**). In *C. elegans*, it is expected that spermatogenic genes are highly expressed and oogenesis genes have reduced expression during the L4 stage, prior to the spermatogenesis-to-oogenesis switch in adulthood. However, the dramatic increase in spermatogenic gene expression and sperm-based sterility in *sosi-1 eri-6[e-f]*Δ mutants suggests the sensors normally fine-tune 22G-RNA levels to restrict spermatogenic transcripts to an appropriate amount of expression.

**Figure 2.**
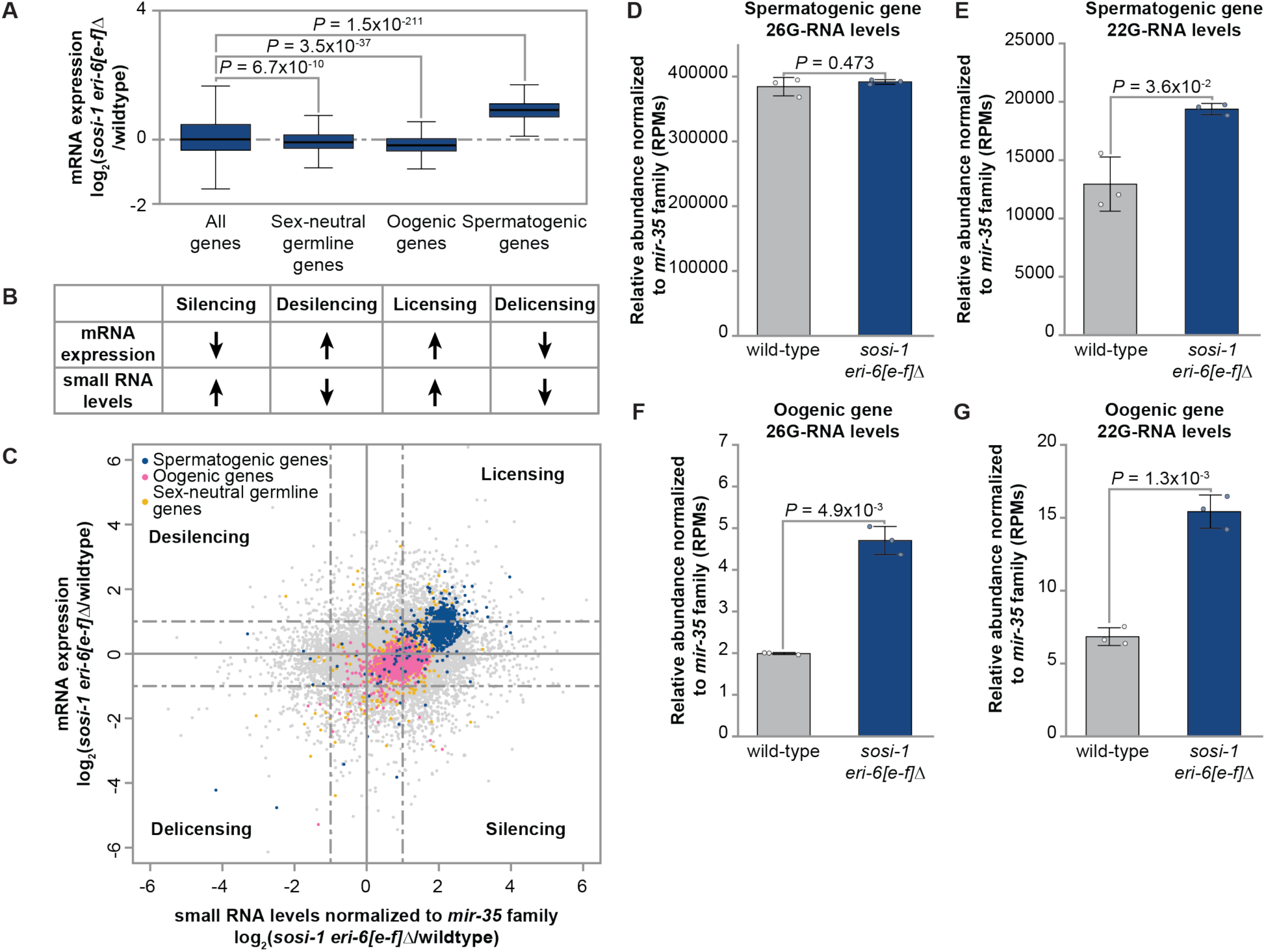
Dysregulation of germline genes in *sosi-1 eri-6[e-f]Δ* mutants. **(A)** Expression changes in *sosi-1 eri-6[e-f]Δ* L4s compared to wild-type L4s for published germline (sex-neutral, oogenic, and spermatogenic) gene lists. Bolded midline indicates median value, box indicates the first and third quartiles, and whiskers represented the most extreme data points within 1.5 times the interquartile range, excluding outliers. Wilcoxon tests were performed to determine statistical significance between the log_2_(fold change) for each gene list compared to all genes, and P-values were adjusted for multiple comparisons. (B) Schema for defining silencing, desilencing, licensing, and delicensing based on increases or decreases in mRNA expression levels and small RNA levels. Up arrows indicate increased levels and down arrows indicated reduced levels. **(C)** Difference in expression (log_2_(fold change)) for genes in *sosi-1 eri-6[e-f]Δ* L4s compared to wild-type animals plotted according to the difference in small **RNA** levels (log_2_(fold change)) normalized to *mir-35* small **RNA** levels in libraries from *sosi-1 eri-6[e-f]Δ* L4s. Each dot represents a gene, with spermatogenic genes (blue), oogenic genes (pink), and sex-neutral germline genes (yellow) highlighted. **(D)** 26G-RNA and (E) 22G-RNA levels normalized to *mir-35* family small **RNA** levels that map to spermatogenic genes are counted, in reads per million (RPMs), for wild-type and *sosi-1 eri-6[e-f]Δ* L4 hermaphrodites. **(F)** 26G-RNA and **(G)** 22G-RNA levels normalized to *mir-35* family small **RNA** levels that map to oogenic genes are counted, in reads per million (RPMs), for wild-type and *sosi-1 eri-6[e-f]Δ* L4 hermaphrodites. Bar graphs represent the mean with dots representing summed RPMs for biological replicates and error bars indicating standard deviation. Two-tailed Welch’s t-tests were performed to determine statistical significance.

We next aimed to determine if changes in small RNA levels correlated with the changes in spermatogenic and oogenic gene expression. If changes in small RNA levels for germline-enriched genes correspond to changes in mRNA levels (**Figure 2B**), it would further establish a causal link between removal of the *sosi-1* and *eri-6[e-f]* sensors and the observed changes in germline gene mRNA levels. To account for differential amplification in each library, we normalized to the relative abundance of small RNAs mapping to members of the *mir-35* family (*mir-35-42*), which are not amplified by the *mutator* complex. Our analyses revealed that spermatogenic genes exhibited increased levels of small RNAs correlating with their increased mRNA levels in *sosi-1 eri-6[e-f]*Δ mutants (**Figure 2C**). When we assessed 26G-RNA and 22G-RNA levels specifically, we found 26G-RNAs mapping to spermatogenic genes were unchanged but 22G-RNAs mapping to spermatogenic genes were significantly increased in *sosi-1 eri-6[e-f]*Δ mutants (**Figures 2D and 2E**). These findings suggest increased 22G-RNA levels aberrantly license the expression of spermatogenic genes in *sosi-1 eri-6[e-f]*Δ L4s. In contrast, we found increased abundance of small RNAs correlated with the down-regulation of oogenic and sex-neutral germline genes in *sosi-1 eri-6[e-f]*Δ mutants (**Figure 2C**). Oogenic genes showed an increase in both 26G- and 22G- RNA levels indicating alterations in oogenic gene targeting by primary and secondary RISCs leads to silencing of oogenic genes in *sosi-1 eri-6[e-f]*Δ L4s (**Figures 2C, 2F, and 2G**). Together, these data indicate the ability to maintain homeostatic 22G-RNA levels through the *sosi-1* and *eri-6[e-f]* sensors is critical for appropriate RNAi-mediated regulation of germline-specific genes during the L4 developmental stage. Furthermore, the siRNA sensors are required to restrict spermatogenic gene expression to a “Goldilocks” zone during spermatogenesis, maintaining homeostasis and fertility.

### Dysregulation of distinct RNAi pathway targets in the absence of *sosi-1* and *eri-6[e-f]* sensors

It is well known that small RNA pathways are critical for germline gene regulation. However, examining the role of small RNA homeostasis in germline gene regulation in animals with fully functional RNAi factors has not been feasible previously. There is overlap between the different RNAi branches that regulate oogenic and spermatogenic transcripts. For example, the *mutator*-dependent ERGO-1 and PRG-1 pathways, as well as the CSR-1 pathway which does not rely on *mutator*-dependent 22G-RNA amplification, have been shown to regulate oogenic transcripts^16,21,22,24,31,32^. Spermatogenic genes are also regulated by the PRG-1 and CSR-1 pathways in addition to the *mutator*-independent ALG-3/4 pathway^25,33-36^. We next assessed changes in genes targeted by ERGO-1, PRG-1, ALG-3/4, and CSR-1 to determine if regulation by these branches is altered upon removal of the *sosi-1 eri-6[e-f]* sensors.

The ERGO-1 pathway functions largely in oocytes and during embryonic development, and not later developmental stages^22,37^. Thus, de-coupling *eri-6/7* expression from the siRNA sensors may have the most impact on ERGO-1 pathway function in oocytes and embryonic developmental stages, while indirect consequences on other RNAi pathways may persist throughout the larval stages. Consistent with this, 26G-RNAs and the mRNA levels for ERGO-1 targets were not significantly changed in *sosi-1 eri-6[e-f]*Δ L4s (**Figures S2A and 3A**), but ERGO-1-target 22G-RNAs were increased (**Figure S2B**). Paired with our previous findings in adult animals^26^, our data indicate that elevated levels of ERGO-1 class 26G-RNAs present in *sosi-1 eri-6[e-f]*Δ embryos trigger increased production of ERGO-1-class 22G-RNAs that persist throughout larval development, but not into adulthood. The dissipation of the ERGO-1-class 22G-RNA levels during the adult stage could point to 1) a lifespan of small RNA populations; 2) a dilution of small RNA levels in the growing germline, or 3) to another molecular mechanism that fine-tunes ERGO-1-class 22G-RNA levels once the animal begins oogenesis.

When we assessed changes in PRG-1, ALG-3/4, and CSR-1 targets, we found that overall PRG-1 and CSR-1 targets were down-regulated and ALG-3/4 targets were up-regulated in *sosi-1 eri-6[e-f]*Δ L4s compared to wild-type animals (**Figures 3A, S3A, and S3B**). The up-regulation of ALG-3/4 targets corresponds to the observed increase in expression of spermatogenic genes, which are largely ALG-3/4 targets (**Figure S3C**). The reduced expression of CSR-1 targets was unexpected as CSR-1 does not rely on the *Mutator* focus for amplification of its 22G-RNAs^16,22,24,38^. However, CSR-1 pathway targets were not significantly enriched amongst the down-regulated genes, suggesting that this overall global change may reflect a subset of CSR-1 targets with substantial change while many CSR-1 targets remain unaffected (**Figure S3B**). We did not expect PRG-1 targets to be overall significantly down-regulated as previous studies had shown in *prg-1* mutants that most piRNA pathway targets are not differentially regulated except for a subset of spermatogenic genes uniquely targeted by the piRNA pathway that become de-silenced during the L4 stage upon loss of complementary *mutator-*dependent 22G-RNAs^39-42^. Assessing changes in PRG-1 targets is further complicated by the fact that some PRG-1 targets are also regulated by CSR-1 and/or ALG-3/4 (**Figures S3C and S3D**). Due to this, we decided to first focus on the unique spermatogenic piRNA targets to determine if they had altered expression in *sosi-1 eri-6[e-f]*Δ L4 animals. Indeed, we observed increased mRNA levels of the uniquely spermatogenic piRNA targets (F26A3.5, ZK930.6, ZK795.2, and Y80D3A.8) in *sosi-1 eri-6[e-f]*Δ L4s compared to wild-type L4s (**Figure 3B**). However, their up-regulation correlated with increased 26G-RNA and 22G-RNA levels (**Figures 3C and 3D**). This was particularly surprising as 21U-RNAs, not 26G-RNAs, are the primary small RNAs that act in the piRNA pathway. When we looked at small RNA levels mapping to all PRG-1 targets, again we observed elevated levels of 22G-RNAs and 26G-RNAs (**Figures S4A and S4B**). Together these findings suggest the potential that a different primary RISC is aberrantly targeting PRG-1 targets, resulting in an increase in their 22G-RNA levels. As ALG-3/4 binds 26G-RNAs and CSR-1 binds *mutator-*independent 22G-RNAs, it is possible the increase in 26G-RNAs and 22G-RNAs mapping to piRNA targets reflect their mis-regulation by ALG-3/4 and CSR-1.

**Figure 3.**
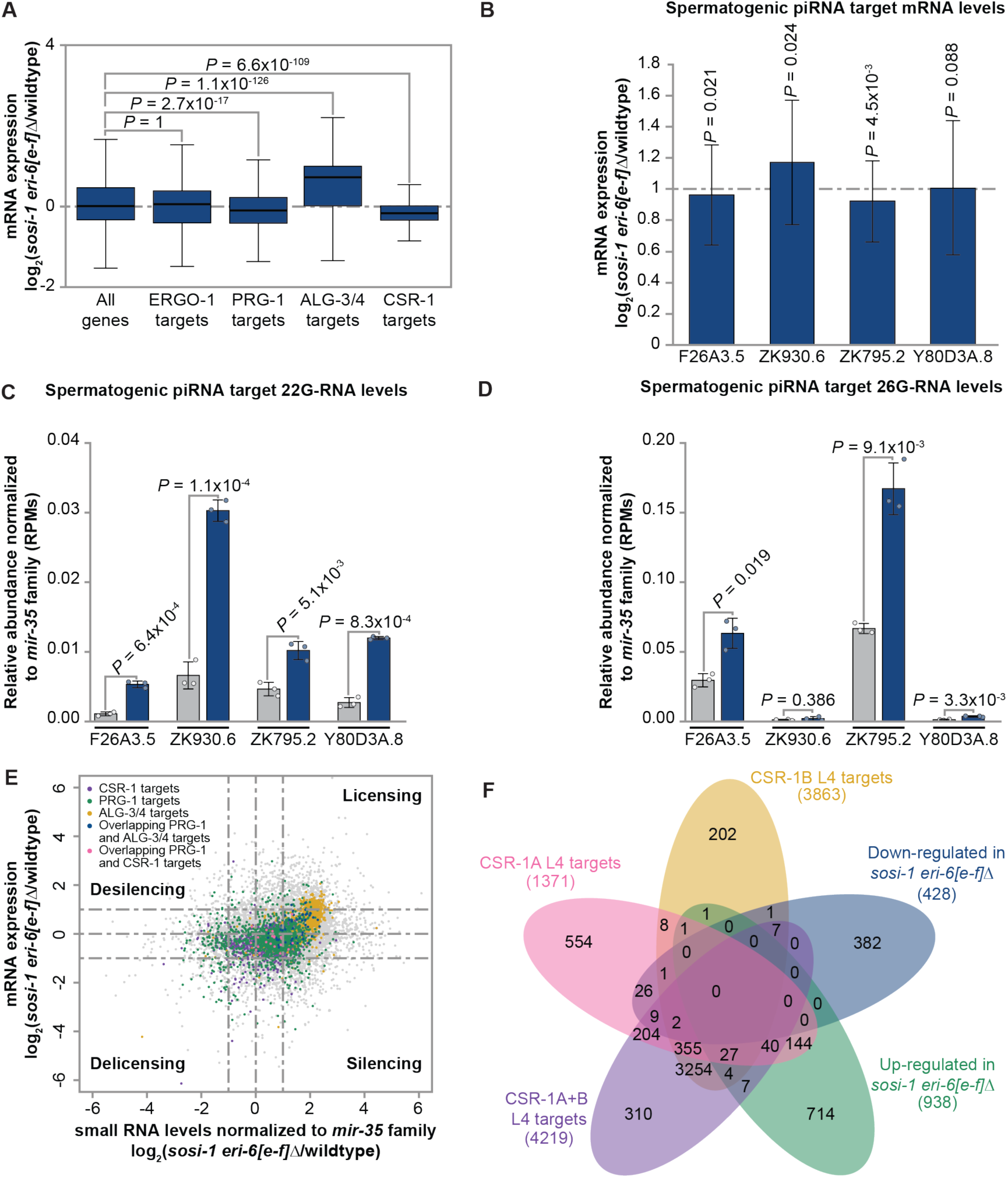
Dysregulation of PRG-1, CSR-1, and ALG-3/4 pathway targets in *sosi-1 eri-6[e-f]Δ* mutants. **(A)** Expression changes in *sosi-1 eri-6[e-f]Δ* L4s compared to wild-type L4s for published ERGO-1, PRG-1, ALG-3/4, and CSR-1 pathway target gene lists. Bolded midline indicates median value, box indicates the first and third quartiles, and whiskers represented the most extreme data points within 1.5 times the interquartile range, excluding outliers. Wilcoxon tests were performed to determine statistical significance between the log_2_(fold change) for each gene list compared to all genes, and P-values were adjusted for multiple comparisons. **(B)** Difference in expression (log_2_(fold change)) for spermatogenic PRG-1 targets in mRNA-seq libraries from *sosi-1 eri-6[e-f]Δ* L4s compared to wild-type animals. Error bars represent log_2_(standard error). P-values were calculated by DESeq2 using the Wald test and corrected for multiple comparisons using the Benjamini and Hochberg method. (C) 22G-RNA and (D) 26G-RNA levels normalized to *mir-35* family small RNA levels that map to spermatogenic piRNA target genes are counted, in reads per million (RPMs), for wild-type and *sosi-1 eri-6[e-f]Δ* L4 hermaphrodites. Bar graphs represent the mean with dots representing summed RPMs for biological replicates and error bars indicating standard deviation. Two-tailed Welch’s t-tests were performed to determine statistical significance. (E) Difference in expression (log_2_(fold change)) for genes in *sosi-1 eri-6[e-f]Δ* L4s compared to wild-type animals plotted according to the difference in small **RNA** levels (log (fold change)) normalized to *mir-35* small **RNA** levels in libraries from *sosi-1 eri-6[e-f]Δ* L4s. Each dot represents a gene, with CSR-1 pathway targets (purple), PRG-1 pathway targets (green), ALG-3/4 pathway targets (gold), overlapping PRG-1 and ALG-3/4 targets (blue), and overlapping PRG-1 and CSR-1 targets (pink) highlighted. **(F)** Venn diagram showing overlap between up-and down-regulated genes in *sosi-1 eri-6[e-f]Δ* L4s with L4-stage targets of CSR-1A, CSR-1B, and CSR-1A+B.

Previous work has shown that loss of piRNA pathway function in *prg-1* mutants leads to misrouting of piRNA targets into the CSR-1 pathway, resulting in their licensing^32^. Further, a previous study demonstrated that direct recruitment of CSR-1 to mature spermatogenic piRNA target transcripts, even in the presence of functional PRG-1, licenses expression of the target^24,32,34,40,41^. It should be noted that *prg-1* and *csr-1* expression is not altered in *sosi-1 eri-6[e-f]*Δ L4 animals, but *alg-3* and *alg-4* mRNA levels are significantly up-regulated in a small RNA-dependent manner in *sosi-1 eri-6[e-f]*Δ L4s (**Figures S4C, S4D, and S4E**). We previously discovered *alg-3* and *alg-4* are regulated by *mutator-*dependent RNAi pathways during the L4 stage to provide thermotolerant sperm-based fertility^43^. Therefore, the excess expression of *alg-3* and *alg-4* is a consequence of the perturbing small RNA levels for distinct RNAi branches caused by removing the ability to fine-tune *mutator-*dependent 22G-RNAs through the sensors. It is possible that removal of the siRNA sensors, and the resulting excess ERGO-1-class input into the *Mutator* focus, initiates a cascade of reduced *mutator* complex amplification of 22G-RNAs for piRNA targets that consequentially leads to their aberrant regulation by other AGOs. We wondered if PRG-1 targets were being distinctly dysregulated based on their co-designation as ALG-3/4 or CSR-1 targets. When we examined mRNA and small RNA levels for genes co-targeted by PRG-1 and ALG-3/4 we found they largely become licensed by increased small RNA targeting in *sosi-1 eri-6[e-f]*Δ animals (**Figure 3E**). In contrast, genes co-targeted by PRG-1 and CSR-1 accumulated small RNAs and became silenced (**Figure 3E**). CSR-1 has two isoforms, CSR-1A and CSR-1B, that act in distinct regions of the L4 germline and regulate spermatogenic and oogenic genes, respectively^34,36^. We found that many of the CSR-1A and CSR-1A+B targets overlapped with genes up-regulated in *sosi-1 eri-6[e-f]*Δ animals (**Figure 3F**). This corresponds with the fact that CSR-1A binds spermatogenic ALG-3/4 targets^36^ and we see increased expression for ALG-3/4 targets and spermatogenic genes (**Figures 2A and 3A**). For the subset of genes down-regulated in *sosi-1 eri-6[e-f]*Δ L4s that overlapped with CSR-1A, CSR-1B, and CSR-1A+B targets, we performed gene ontology enrichment analyses and found they were enriched for structural constituents of chromatin (**Figure S4F**). Therefore, the down-regulation observed when assessing all annotated CSR-1 targets (**Figure 3A**) reflects significantly reduced expression of some unique CSR-1 targets and PRG-1/CSR-1 co-regulated targets while spermatogenic CSR-1 targets are up-regulated. Taken together, our analyses reveal a distinct change in fate (silencing, desilencing, licensing, or delicensing) for targets when 22G-RNA homeostasis is disrupted based on their co-regulation by the piRNA pathway and ALG-3/4 or CSR-1. Our findings indicate sensor-mediated homeostasis of *mutator-*dependent 22G-RNAs is required to prevent incorrect sorting of germline gene targets between the piRNA, CSR-1, and ALG-3/4 pathways. Further, the observed perturbations in ALG-3/4 and CSR-1 target regulation clearly indicate that impairing the sensor-mediated feedback has severe consequences that extend across multiple RNAi branches, including ones that do not utilize the *mutator* complex.

### Sensor-mediated 22G-RNA homeostasis is required for histone gene expression

Our observation that CSR-1-targeted structural constituents of chromatin are down-regulated led us to examine histone gene expression to determine if changes in CSR-1 targeting or piRNA pathway disfunction is occurring in *sosi-1 eri-6[e-f]*Δ L4 animals. Recent studies have shown that histone gene transcripts are down-regulated in both *prg-1* and *csr-1* mutants, but the mechanism underlying reduced histone expression in each of these mutants differs^39,40,44^. CSR-1 and its associated 22G-RNAs are required for 3’ end cleavage of histone pre-mRNAs as part of their processing into mature histone mRNAs^44^. In animals lacking functional CSR-1 or its 22G-RNAs, histone pre-mRNAs accumulate without being processed, resulting in overall reduced histone expression^44^. In contrast, loss of functional PRG-1 causes the AGO WAGO-1 to delocalize from perinuclear germ granules and aberrantly associate with CSR-1 in the cytoplasm^40^. This interaction results in increased CSR-1-derived 22G-RNAs for histone transcripts being loaded into WAGO-1 and initiating histone transcript silencing^39,40^. We hypothesized that if removal of the sensors directly impairs *mutator-*dependent production of piRNA-target 22G-RNAs, then in their absence, WAGO-1 could associate with CSR-1 leading to an increase in 22G-RNAs mapping to the histone genes and reduced histone expression. As *prg-1* and *csr-1* expression is not significantly altered in *sosi-1 eri-6[e-f]*Δ L4 animals, changes in histone gene mRNA and small RNA levels would indicate aberrant routing of these targets due to pathway dysfunction driven by the inability to fine-tune *mutator-*dependent amplification of 22G-RNAs via the sensors (**Figure S4C**). Indeed, compared to wild-type L4 animals, *sosi-1 eri-6[e-f]*Δ mutants had significantly reduced histone gene expression (**Figures 4A, 4B, 4C, and S5A**). When assessed individually, members of each replication-dependent histone family (H2A, H2B, H3, and H4) had significantly reduced expression (**Figure 4C**). In contrast, most members of the replication-independent histone family (H3.3) did not change significantly, with the exception of the spermatogenesis-specific histone variant, *his-70*^45^, which was up-regulated (**Figure 4C**).

**Figure 4.**
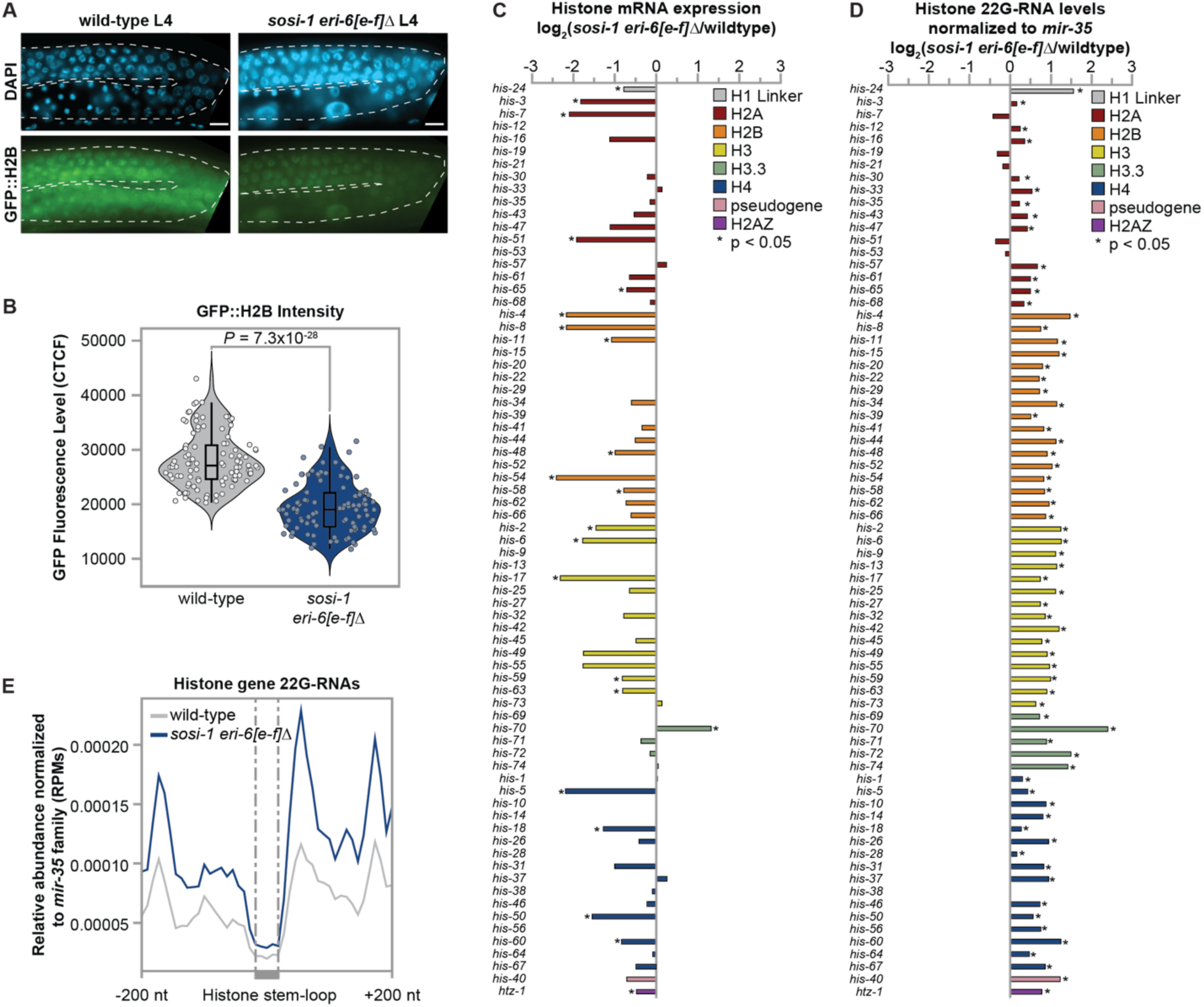
Sensor-mediated 22G-RNA homeostasis is required for histone gene expression. **(A)** Representative innages of germlines in wild-type and *sosi-1 eri-6[e-f]Δ* L4 animals expressing GFP::H2B. Scale bar indicates 10pm. **(B)** Violin plots showing qunatification of GFP::H2B expression in pachytene germ cells in corrected total cell fluorescence (CTCF). Each dot represents a pachytene germ cell, with 10 cells assessed from each of 10 germlines. Bar plots displaying log_2_(fold change) of **(C)** mRNA levels and **(D)** 22G-RNA levels normalized to *mir-35* family small RNA levels for each histone gene in *sosi-1 eri-6[e-f]Δ* L4s compared to wild-type L4s. Bars are colored by histone families and * indicates a P-value less than 0.05 was calculated and corrected for multiple comparisons using the Benjamini and Hochberg method. **(E)** Metaprofile analysis showing the distribution of 22G-RNAs normalized to the *mir-35* family (RPMs) 200 nucleotides up and down stream of the stem-loop sequences of histone genes in wild-type (gray) and *sosi-1 eri-6[e-f]Δ* (blue) L4 animals. The average of three biological replicates is shown.

Our bioinformatic analyses also revealed that 22G-RNA levels for each histone gene were increased in *sosi-1 eri-6[e-f]*Δ mutants (**Figures 4D, S5B and S5C**). Similarly to what was observed in *prg-1* mutants^40^, meta-profile analyses showed in *sosi-1 eri-6[e-f]*Δ mutants there is an increase in 22G-RNAs mapping to histone gene bodies, with a bias towards the end of histone genes before the stem-loop structure that is bound by CDL-1 and CSR-1^46^ (**Figures 4E and S5C**). Interestingly, we also see 22G-RNAs downstream of the stem-loop (**Figure 4E)**, suggesting CSR-1 is processing the full length of the histone mRNAs into 22G-RNAs that can be loaded into WAGO-1. A similar spreading of histone mRNA processing was observed in *prg-1* mutant young adults upon knockdown of CDL-1, which helps direct CSR-1 activity to the stem-loop structure of histone transcripts^40^. In *C. elegans,* CDL-1 has strong nuclear localization in somatic cells, embryos, and maturing oocytes and is required maternally for histone mRNA maturation and fertility^44,47,48^. However, whether CSR-1 and CDL-1 interact during the L4 developmental stage when spermatogenesis is occurring remains unclear as previous studies examined their interaction during the adult stage. In *sosi-1 eri-6[e-f]*Δ L4 mutants, *cdl-1* expression was not altered at the mRNA level and wild-type L4 animals also exhibited 22G-RNA production downstream of the stem-loop, so it is possible that the spreading of 22G-RNAs along the histone transcripts could reflect that CDL-1 does not focus CSR-1 processing of histone transcripts during the L4 stage (**Figures S5D and 4E**). Together, these results further support that removal of the *sosi-1* and *eri-6[e-f]* sensors directly impacts the function of the piRNA pathway downstream of the *mutator* complex and demonstrates that the sensors’ roles in balancing *mutator-dependent* 22G-RNA classes impacts the regulation of the replication dependent histone genes required for proper fertility and development.

### Loss of sensor-mediated 22G-RNA homeostasis leads to defects in pachytene germ cells

All our sequencing analyses indicate that loss of the *sosi-1* and *eri-6[e-f]* sensors regulating *eri-6/7* triggers a cascade that results in dysfunction of multiple RNAi pathways regulating germline and histone genes. We next looked to see if these molecular changes impacted germline development during spermatogenesis. The *C. elegans* germline develops from a pool of mitotically replicating nuclei at the distal end that travel in a gradient as they undergo meiosis, such that germ cells in the earlier stages of meiosis are closer to the distal ends than germ cells in later meiotic stages^49^. This allows for the number of germ cells in each stage of meiosis to be quantified. At initial glance, the organization, overall morphology, and size of *sosi-1 eri-6[e-f]*Δ L4 hermaphroditic and male germlines resembled those of wild-type animals. Further, the integrity and morphology of P granules, perinuclear germ granules adjacent to *Mutator* foci and a hall mark of germ cell identity, remained intact in *sosi-1 eri-6[e-f]*Δ L4s (**Figure 5A**). However, we observed an increased incidence of males in the *sosi-1 eri-6[e-f]*Δ populations (not quantified) that suggested *sosi-1 eri-6[e-f]*Δ animals have defects in meiosis. Male *C. elegans* arise due to meiotic chromosome non-disjunction events^50^ and can be propagated in a population through mating. As *sosi-1 eri-6[e-f]*Δ mutants are sperm-based sterile, the increased number of males found in the population is indicative of higher rates of meiotic chromosome non-disjunction than in wild-type populations. We decided to quantify the number of meiotic germ cells, and found that *sosi-1 eri-6[e-f]*Δ males and L4s had significantly reduced numbers of meiotic germ cells compared to wild-type animals (**Figures 5B and 5C**). When we counted the number of meiotic germ cells specifically in the transition and pachytene zones, we found that *sosi-1 eri-6[e-f]*Δ mutant L4s and males had fewer pachytene germ cells than their wild-type counterparts (**Figures 5D and 5E**). This indicates the molecular changes triggered by removal of the siRNA sensors leads to germline defects in the pachytene zone, which is critical for spermatogenesis.

**Figure 5.**
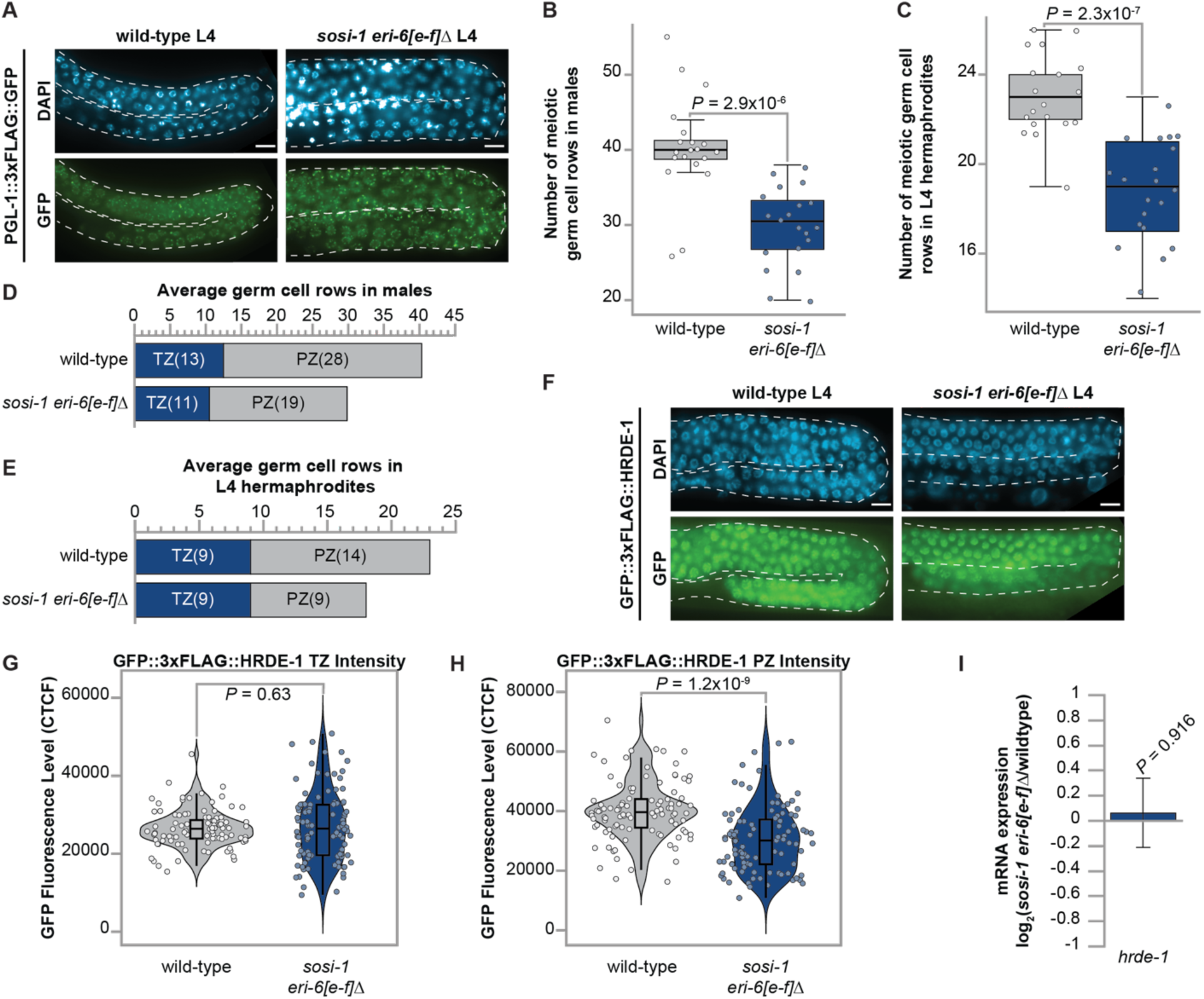
Removal of 22G-RNA sensors leads to pachytene defects in the germline of males and L4 hermaphrodites. **(A)** Representative images of germlines in wild-type and *sosi-1 eri-6[e-f]Δ* L4 animals expressing PGL-1::3xFLAG::GFP. Scale bar indicates 10pm. Box plots depicting total number of meiotic germ cell rows in wild-type and *sosi-1 eri-6[e-f]Δ* **(B)** males and **(C)** L4 hermaphrodite animals. For (B and C) circles represent individual germlines. Bolded midline indicates median value, box indicates the first and third quartiles, and whiskers represented the most extreme data points within 1.5 times the interquartile range, excluding outliers. N = 20. Number of germ cell rows in wild-type and *sosi-1 eri-6[e-f]Δ* **(D)** male and **(E)** L4 hermaphrodite animals showing average number of cells in the transition zone (TZ, blue) and pachytene zone (PZ, gray). The average of 20 individual germlines is shown. **(F)** Representative images of germlines in wild-type and *sosi-1 eri-6[e-f]Δ* L4 animals expressing GFP::3xFLAG::HRDE-1. Scale bar indicates 10pm. Violin plots showing quantification of GFP::3xFLAG::HRDE-1 levels in **(G)** transition zone and **(H)** pachytene zone germ cells in corrected total cell fluorescence (CTCF). Each dot represents a germ cell, with 10 cells assessed from each of 10 germlines. **(I)** Expression changes in *sosi-1 eri-6[e-f]Δ* L4s compared to wild-type animals for *hrde-1.* Error bars indicate log_2_(standard error) and P-value was calculated and corrected for multiple comparisons using the Benjamini and Hochberg method. For **(B, C, G, and H)** statistical significance was calculated by two-tailed Welch’s t test, mutants were compared to wild-type.

Pachytene piRNAs have been shown to be essential for spermatogenesis across many animals^51,52^. In *C. elegans,* a previous study found that *prg-1* mutant males also had shortened pachytene zones due to defects in meiotic progression^42^. Further, another recent study demonstrated that piRNA pathway function is critical specifically within pachytene nuclei of L4 germlines for localization of the nuclear AGO, HRDE-1, which binds *mutator-*dependent 22G-RNAs for PRG-1-targeted spermatogenic transcripts^41^. The delocalization of HRDE-1 in *prg-1* mutants also coincided with increased L4-stage expression of spermatogenic transcripts, similar to what we observe in *sosi-1 eri-6[e-f]*Δ L4s (**Figure 2A**). Our bioinformatic analyses revealed spermatogenic transcripts targeted by PRG-1, ALG-3/4, and CSR-1 are dysregulated in *sosi-1 eri-6[e-f]*Δ L4s, therefore, we suspected there may be a dearth of *mutator-*dependent 22G-RNAs complementary to these transcripts for HRDE-1 to bind, resulting in loss of nuclear localization of HRDE-1 in pachytene germ cells. To test this hypothesis, we quantified the GFP::3xFLAG::HRDE-1 nuclear localization within transition zone and pachytene zone nuclei from the germlines of wild-type and *sosi-1 eri-6[e-f]*Δ mutant L4 animals. Indeed, transition zone nuclei did not have changes in HRDE-1 localization, but pachytene germ cells had a significant reduction of nuclear HRDE-1 localization despite no change in *hrde-1* expression (**Figures 5F, 5G, 5H and 5I**). These findings, taken together with the alterations in mRNA and small RNA levels for spermatogenic genes, demonstrate that the 22G-RNA sensors are critical for coordinating key developmental programs in the pachytene zone of the germline during spermatogenesis.

### Spermiogenesis defects underly the sperm-based sterility in *sosi-1 eri-6[e-f]*Δ mutants

The observed unfertilized oocytes from hermaphrodites and failure of males to produce progeny upon mating indicates that the reproductive potential of both hermaphrodite- and male-derived sperm of *sosi-1 eri-6[e-f]*Δ mutants is negatively impacted (**Figures 1C, 1D, and 1E**). Loss of sperm-based fertility could be triggered by defects in spermatid production during spermatogenesis, spermatid activation during spermiogenesis, or in the sperm’s ability to complete fertilization. We first looked at the expression levels of genes critical for successful completion of spermatogenesis, spermiogenesis, and fertilization to see if changes at the mRNA level would point to which process was affected in *sosi-1 eri-6[e-f]*Δ L4 hermaphrodites. Fitting with the observation that as a class sperm-enriched genes are up-regulated, some of these essential genes were up-regulated (*spe-4, spe-6, spe-26, spe-10, spe-17, fer-1, spe-27,* and *spe-11*) while the rest showed no significant change in expression (**Figure 6A**). To determine if defects in spermatogenesis triggered the loss of sperm-based fertility, we DAPI-stained wild-type and *sosi-1 eri-6[e-f]*Δ mutant animals and assessed the abundance of post-meiotic spermatids present in the spermatheca. We found that on average, *sosi-1 eri-6[e-f]*Δ mutant males had significantly more spermatids than their wild-type counterparts (**Figure 6B**). This finding, paired with our observation that *sosi-1 eri-6[e-f]*Δ mutant males have fewer pachytene germ cells than wild-type males, suggests germ cells are transitioning through the pachytene zone more rapidly due to defects in meiotic progression (**Figure 5D**). We next examined *sosi-1 eri-6[e-f]*Δ mutant hermaphrodites at the very beginning of the young adult stage, after spermatogenesis has completed but prior to oocytes being fertilized to see if the change in spermatid abundance occurred in both sexes. Interestingly, *sosi-1 eri-6[e-f]*Δ mutant hermaphrodites had fewer post-meiotic spermatids than their wild-type counterparts (**Figure 6C**). The sex-specific differences in number of post-meiotic spermatids, despite both *sosi-1 eri-6[e-f]*Δ mutant males and L4 hermaphrodites having fewer pachytene germ cells than their wild-type counterparts, may reflect sex-specific differences in the rate of progression through the meiotic stages or in the total number of germ cells that underwent spermatogenesis. However, the fact that *sosi-1 eri-6[e-f]*Δ mutant males and hermaphrodites produce sufficient numbers of post-meiotic spermatids indicates that defects in spermatogenesis do not drive the loss of sperm-based fertility.

**Figure 6.**
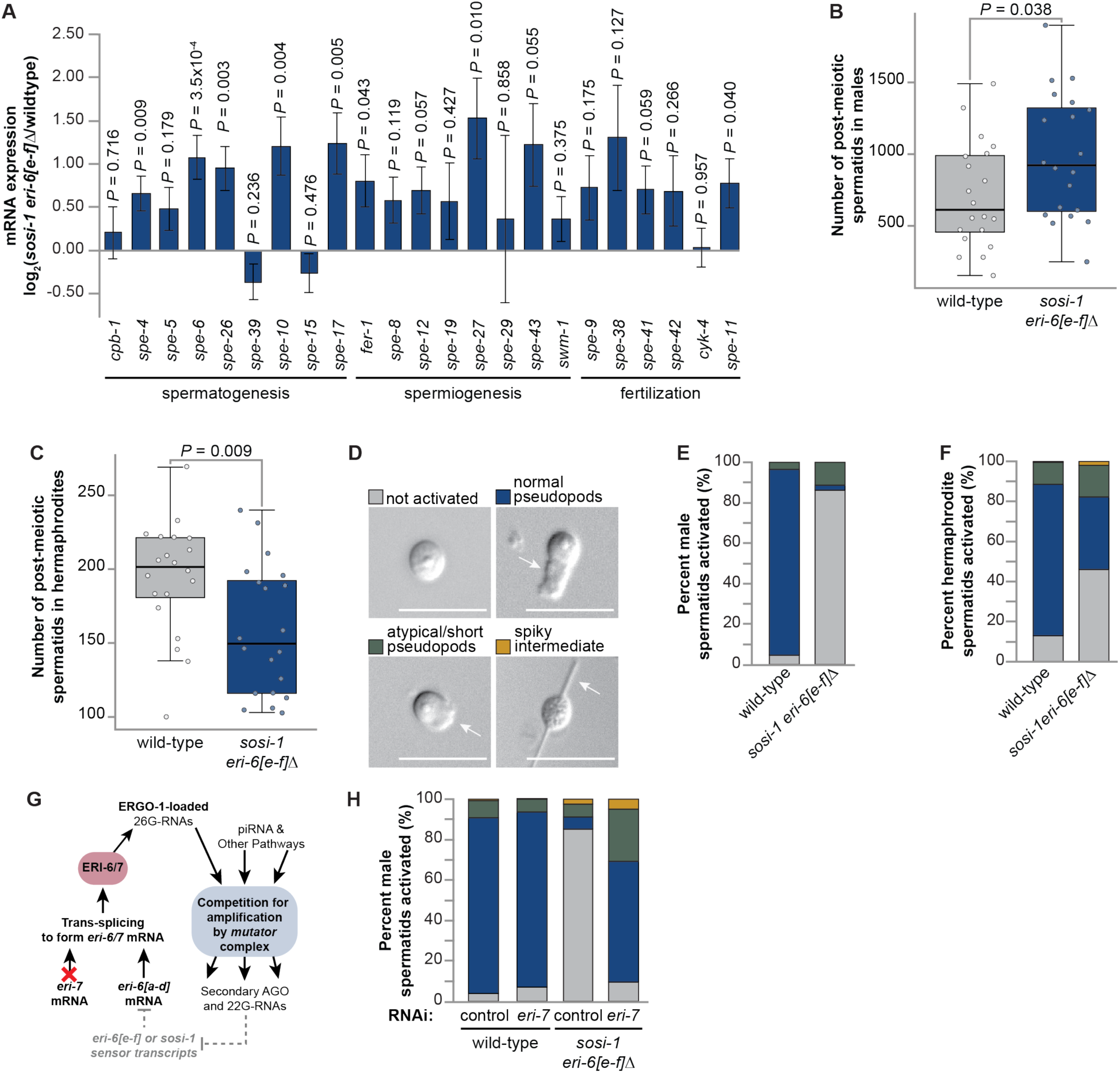
Removal of 22G-RNA sensor triggers spermiogenesis defects. **(A)** Difference in expression (log_2_(fold change)) for genes critical for spermatogenesis, spermiogenesis, and fertilization in mRNA-seq libraries from *sosi-1 eri-6[e-f]Δ* L4s compared to wild-type animals. Error bars represent log_2_(standard error). P-values were calculated by DESeq2 using the Wald test and corrected for multiple comparisons using the Benjamini and Hochberg method. Box plots depicting total number of post-meiotic spermatids for wild-type (gray) and *sosi-1 eri-6[e-f]Δ* (blue) **(B)** males and **(C)** L4 hermaphrodites. Circles represent individual animals. Bolded midline indicates median value, box indicates the first and third quartiles, and whiskers represented the most extreme data points within 1.5 times the interquartile range, excluding outliers. N = 20. For **(B and C)** statistical significance was calculated by two-tailed Welch’s *t* test, mutants were compared to wild-type. **(D)** Representative images of spermatids after exposure to Pronase E that remain unactivated (gray), develop normal psuedopods (blue), develop atypical pseudopods (green), or arrest as spiky intermediates (gold). Scale bar indicates 10pm and white arrows point to pseudopods. Percent of Pronase E-treated spermatids that are not activated (gray), have normal pseudopod formation (blue), or atypical pseudopod formation (green), or arrest as spiky intermediates (gold) are shown for wild-type and *sosi-1 eri-6[e-f]Δ* **(E)** adult males and **(F)** young adult hermaphrodites. **(G)** Schema of the feedback mechanism including *eri-6[a-d]* and *eri-7* trans-splicing to from functional *eri-6/7* mRNAs. Gray text and gray dashed lines show interactions that are ablated in *sosi-1 eri-6[e-f]Δ* mutants. Red X shows RNAi of *eri-7* will reduce *eri-7* mRNA levels, leading to reduced trans-splicing of functional *eri-6/7* mRNAs. **(H)** Percent of Pronase E-treated spermatids that are not activated (gray), have normal pseudopod formation (blue), or atypical pseudopod formation (green), or arrest as spiky intermediates (gold) are shown for wild-type and *sosi-1 eri-6[e-f]Δ* adult males fed bacteria producing control (L4440) dsRNA or dsRNA for *eri-7.* For **(E, F, and H)** N ≥ 200 spermatids per genotype per condition.

Non-motile spermatids develop pseudopods and become motile spermatozoa during spermiogenesis after exposure to an activating agent^53^. While we did not observe reduced expression of genes critical for spermiogenesis (**Figure 6A**), it has been previously shown that spermatids of *C. elegans* mutants with overexpression of sperm-enriched genes fail to activate^41^. To determine if spermiogenesis defects underly the loss of sperm reproductive potential, we performed an *in vitro* activation assay on spermatids dissected from wild-type and *sosi-1 eri-6[e-f]*Δ mutant males and hermaphrodites. After exposure to Pronase E, we assessed pseudopod morphology and classified each spermatozoan as not activated, normal, atypical, or as spiky intermediates (**Figure 6D**). Strikingly, 91.74% of wild-type male spermatids formed normal pseudopods whereas only 2.34% of *sosi-1 eri-6[e-f]*Δ mutant male spermatids formed normal pseudopods and 86.33% failed to activate (**Figure 6E**). When we assessed hermaphrodite-derived spermatids, we found sex-specific rates of activation in *sosi-1 eri-6[e-f]*Δ mutants. 75.71% of wild-type and 36.06% of *sosi-1 eri-6[e-f]*Δ mutant hermaphrodite-derived spermatids formed normal pseudopods after exposure to Pronase E (**Figure 6F**). The ability for some hermaphrodite-derived spermatids to properly activate but complete failure for male-derived spermatids to activate in *sosi-1 eri-6[e-f]*Δ mutants likely reflects differences in male and hermaphrodite sperm activation pathways^53-56^. Furthermore, the sperm activation assay results corroborate our findings where *sosi-1 eri-6[e-f]*Δ mutant hermaphrodites are able to self-fertilize and produce progeny, albeit at a reduced number compared to wild-type animals, and *sosi-1 eri-6[e-f]*Δ mutant males are unable to produce progeny upon mating (**Figures 1B and 1E**). Overall, we found loss of the ability to fine-tune *mutator*-dependent 22G-RNA levels is critical for spermiogenesis in both *C. elegans* sexes. Our findings highlight perturbations to RNAi homeostasis triggers sex-specific differences in the penetrance of spermiogenesis defects and indicates that male-derived spermatids are more sensitive to perturbations than hermaphrodite-derived spermatids.

If the spermiogenesis defects are a direct consequence of un-regulated production of ERI-6/7 upon removal of the siRNA sensors, leading to ERGO-1-class 22G-RNA production overwhelming *mutator* complex resources, we would expect that reducing ERI-6/7 levels in the *sosi-1 eri-6[e-f]*Δ mutants would rescue the spermiogenesis defects (**Figure 6G**). To test this, we performed an RNAi feeding assay to knockdown *eri-7* transcripts that must be trans-spliced to the *eri-6[a-d]* isoforms to create functional *eri-6/7* mRNA. Then we assessed the sperm activation of male-derived spermatids isolated from wild-type and *sosi-1 eri-6[e-f]*Δ mutants fed the *eri-7* RNAi clone or a control RNAi clone (L4440). Spermatid activation rates in wild-type males exposed to the control and *eri-7* RNAi clones, and *sosi-1 eri-6[e-f]*Δ males exposed to the control RNAi clone were similar to the spermatid activation rates observed in males not exposed to RNAi clones (**Figures 6E and 6H**). Strikingly, the rate of spermatids forming normal pseudopods in *sosi-1 eri-6[e-f]*Δ males increased from 5.67% in animals exposed to the RNAi control to 59.67% in animals exposed to the *eri-7* RNAi clone (**Figure 6H**). This result further supports our model and demonstrates that fine-tuning of ERI-6/7 levels, and thus ERGO-1-class 26G-RNA levels that feed into the *Mutator* focus, is critical for maintaining appropriate RNAi-mediated regulation required for physiological processes such as sperm development, and ultimately fertility.

## DISCUSSION

All previous studies that have shown disrupting RNAi-mediated gene regulation leads to germline gene dysregulation and loss of fertility across species have relied on mutating key RNAi pathway factors, thus rendering at least one RNAi branch completely non-functional. Our *sosi-1 eri-6[e-f]*Δ mutants are the first mutant in any species that has intact RNAi pathway factors but lack the ability to maintain small RNA homeostasis. Here we leveraged this unique mutant to examine the molecular and physiological consequences of disrupting small RNA homeostasis in the presence of all functional RNAi factors. Supporting our model that the *sosi-1* and *eri-6[e-f]* sensors are critical for allowing dynamic balancing of *mutator* complex amplified 22G-RNAs, we found that removal of the siRNA sensors, and thus the loss of regulatory potential on ERI-6/7 levels, leads to increased abundance of ERGO-1-class 22G-RNAs during the larval stage when spermatogenesis occurs in *C. elegans*. Animals lacking the siRNA sensors exhibited a loss of sperm-based fertility. Thus, sperm development and reproductive potential are sensitive to disruptions to homeostatic small RNA levels.

The loss of sperm-based fertility in *sosi-1 eri-6[e-f]*Δ mutants coincided with increased expression of spermatogenesis genes during spermatogenesis, indicating that levels of sperm-enriched transcripts need to be restricted to a “Goldilocks zone” of expression and that if they exceed a threshold, it can negatively impact sperm development. During the L4 developmental stage when *C. elegans* are undergoing spermatogenesis, sperm-enriched transcripts are regulated by several RNAi branches to fine-tune their expression levels. Many of the previous studies focused on understanding RNAi pathway branches that coordinate germline gene expression to restrict spermatogenesis to the L4 developmental stage in *C. elegans* or help to coordinate the spermatogenesis-to-oogenesis switch. For example, the primary AGOs ALG-3 and ALG-4 have been shown to positively and negatively regulate sperm transcripts, and downstream CSR-1 binds 22G-RNAs amplified from the ALG-3/4 targets to further fine-tune their expression and ensure temporal precision when the germline switches from spermatogenesis to oogenesis^33,34,36,57^. While the ALG-3/4 and CSR-1 pathways do not rely on the *mutator* complex for amplification of 22G-RNAs, both pathways are critical during the L4 stage and are intimately linked to *mutator-*dependent RNAi pathways. Our previous work discovered an RNAi-to-RNAi regulatory cascade is responsible for maintaining appropriate expression of *alg-3* and *alg-4* to restrict ALG-3/4 pathway function and the spermatogenesis developmental program to the L4 stage^43^. This RNAi-to-RNAi cascade is particularly important for sperm reproductive potential during heat stress. We also observed aberrant regulation of *alg-3* and *alg-4* in *sosi-1 eri-6[e-f]*Δ mutants, which are experiencing a form of molecular stress due to their inability to maintain small RNA homeostasis. We believe the disruption to RNAi homeostasis upon removal of the *sosi-1* and *eri-6[e-f]* sensors triggers a series of consequences within RNAi pathways that prevents the *mutator*-dependent cascade that regulates *alg-3/4* from properly functioning. The dysregulation of *alg-3/4* expression, and thus ALG-3/4 pathway function, would contribute to the observed up-regulation of spermatogenic transcripts. This finding further reveals the interdependences between the distinct RNAi branches and highlights the importance of uncovering the regulatory architecture embedded within RNAi pathways to improve our understanding of how these regulatory pathways function.

The piRNA pathway plays essential roles in animal germ cells to maintain both oocyte-based and sperm-based fertility by repressing transposable elements, and as more recently appreciated, in regulating some spermatogenic protein-coding genes^28,35,58^. Indeed, aberrant increases in sperm-enriched transcripts during spermatogenesis, similar to what we found in *sosi-1 eri-6[e-f]*Δ mutants, have been observed in animals lacking the Piwi-clade AGO PRG-1 or HRDE-1, which loads downstream piRNA-target 22G-RNAs^7,35,41^. An antagonistic behavior in balancing sperm-enriched transcripts is evident between the CSR-1 and piRNA pathways, as loss of functional PRG-1 or CSR-1 causes misrouting of the sperm transcripts into the other pathway, resulting in their dysregulation^24,32,34,40,41^. However, even with functional AGOs present, misrouting can occur, as shown when direct recruitment of CSR-1 to mature spermatogenic piRNA target transcripts in the presence of functional PRG-1 resulted in licensing expression of the target^41^. Our mRNA and small RNA sequencing results suggest that the small RNA populations, and thus AGO-loading, targeting spermatogenetic transcripts are altered when the siRNA sensors are removed and small RNA homeostasis cannot be maintained. Importantly, the expression of CSR-1, PRG-1, HRDE-1, and formation of P granules were not altered in *sosi-1 eri-6[e-f]*Δ mutants. This indicates that the requirement to balance *mutator-*dependent 22G-RNAs supersedes the presence of functional AGOs and P granules. Furthermore, our study has demonstrated that disrupting RNAi homeostasis in the presence of functional AGOs and germ granules has catastrophic effects on a broad range of physiological processes. For example, we uncovered that *sosi-1 eri-6[e-f]*Δ mutants have inappropriate silencing of histone genes, triggering reduced histone incorporation in the germline. We also show *sosi-1 eri-6[e-f]*Δ mutants have shortened pachytene region indicative of defects in the rate of meiotic progression. On their own, defects in histone incorporation into chromatin and chromatin compaction or loss of meiotic regulation leading to germ cells proceeding through meiosis without clearing important checkpoints, can cause germ cells to lose their reproductive potential. Our results highlight the role not only RNAi pathways, but homeostasis of small RNA levels, play in coordinating meiosis and maintaining germ cell integrity.

Taken together, our bioinformatic analyses and cytological observations indicate that target regulation by the piRNA pathway is hyper-sensitive to perturbations in dynamic balancing of *mutator* complex resources in the germline. It should be noted, the *mutator* complex is not fully monopolized by ERGO-1-class 22G-RNAs in *sosi-1 eri-6[e-f]*Δ mutants and PRG-1-recognized targets still have the potential to have 22G-RNAs amplified in the *Mutator* focus. Thus, our physiological assays reveal the importance of the subtle ability to intentionally shift production of distinct 22G-RNA classes by the *mutator* complex based on cellular needs, developmental context, and environmental factors. While *sosi-1 eri-6[e-f]*Δ mutants produced abundant numbers of post-meiotic spermatids, the physiological consequence of disrupting RNAi homeostasis was a failure for the spermatids to successfully execute spermiogenesis. Loss of functional PRG-1 or HRDE-1 also leads to spermiogenesis defects, although to a less severe degree than we observed in *sosi-1 eri-6[e-f]*Δ mutant males^41^. This indicates that despite all RNAi pathway factors being present and functional, disrupting the ability to fine-tune the input levels for the *mutator* complex has catastrophic effects on the piRNA pathway’s ability to maintain appropriate target regulation. Studies of piRNA pathway mutants in different organisms have shown the piRNA pathway is critical for spermatogenesis, and arrest in spermatogenesis has been observed in mice with mutations in the PIWI-homologs, MIWI and MILI^59,60^. Furthermore, mice lacking the ability to make full piRNA precursors from the pi6 piRNA cluster were shown to have increased sperm-enriched gene expression and spermiogenesis defects^61^ matching what we observe in *sosi-1 eri-6[e-f]*Δ mutants. Our findings in *sosi-1 eri-6[e-f]*Δ mutants reveal that the presence of PRG-1, piRNA precursors, and other piRNA pathway upstream machinery is not sufficient to maintain proper piRNA-mediated regulation of sperm genes in the absence of the siRNA sensors’ ability to act like a rheostat for ERGO-1 targets inputted into the *Mutator* focus.

If removal of the siRNA sensors triggers molecular perturbations to the piRNA pathway in *C. elegans,* is the piRNA pathway function sensitive to fine-tuning by molecular feedback motifs in other organisms? To date, few studies have focused on identifying regulatory motifs embedded within small RNA pathway networks, thus there is exciting potential for more mechanisms that maintain small RNA homeostasis to be uncovered. A recent study found the *sosi-1* and *eri-6[e-f]* sensors regulatory control over the trans-spliced *eri-6/7* transcript likely originated from the arms race between transposons and small RNA pathways in *C. elegans*^62^. The structural rearrangement of the *eri-6-7* gene, *sosi-1*, and what is annotated as *eri-6[e-f]* were induced by multiple insertions of the DNA transposon, *Polinton*^62^. This likely led to potentiated function of the ERGO-1 pathway, which regulates retrotransposons, freeing up *Mutator* resources for 22G-RNA production for piRNA and exogenous RNAi targets. However, transposons are a threat to genome integrity within the germline, and thus the ability to maintain ERGO-1 pathway function would be necessary. Further, *eri-6[f]*, which shares 97.3% sequence identity with another gene of unknown function, K09B11.4, is predicted by protein BLAST to have potentially originated from a *gypsy* retrotransposon^26,62^. This suggests the siRNA sensors’ genomic structure not only originated due to transposon activity, but also that *eri-6[f]* ‘s function as an siRNA sensor could have arisen through co-option of a *gypsy* transposon by the RNAi pathways. The co-option of transposons by host genomes has occurred with a common frequency across evolution and has been observed to re-structure gene regulatory networks in somatic tissues^63-65^. A recent study found that transposon movement in the genome of *Caenorhabditis* nematodes had led to extensive co-option of transposons in the regulation of germline genes^66^. Further, another study revealed the piRNA pathway in chickens has been co-opted to regulate a transposon-derived gene that temporally regulates specification of the neural crest during development^67^. These studies are part of the growing body of evidence that transposons can be co-opted and regulated by RNAi pathways to coordinate the execution of distinct and varied physiological processes. It is possible natural selection favored the evolution of the *sosi-1* and *eri-6[e-f]* sensors as a molecular rheostat for the ERGO-1-class input into the *Mutator* focus to balance competing needs between regulating protein-coding genes required for fertility and repressing transposons within the germline while integrating environmental and organismal cues. Taken together, our results demonstrate the ability to maintain homeostatic balance between distinct classes of small RNAs is critical for ensuring robust RNAi-meditated gene regulation in the germline and has profound and wide-ranging roles in diverse physiological processes.

## DATA AVAILABILITY

High-throughput sequencing data for mRNA-seq and small RNA-seq experiments generated during this study are available through Gene Expression Omnibus (GEO #: GSE324389 and GSE324390). Source data are provided with this paper.

## ACKNOWLEDGEMENTS

We thank the members of the Rogers lab for helpful discussions. Next generation sequencing was performed by Zibiao Guo, who we thank for his technical support, at the North Texas Genome Center.

*Author contributions:* Conceptualization, A.K.R.; Investigation, H.M. and A.K.R; Formal Analysis, H.M. and A.K.R; Writing – original draft, H.M.; Writing – reviewing and editing, H.M. and A.K.R; Supervision, A.K.R; Funding Acquisition, A.K.R.

## MATERIALS AND METHODS

### C. elegans strains

Unless otherwise stated, worms were grown at 20⁰C on NGM plates seeded with OP50-1 *Escherichia coli* according to standard conditions^68^. All strains are in the N2 background. Strains used include:

- N2 – wild-type.
- BS553 – *fog-2(oz40) V*.
- USC1338 – *sosi-1 eri-6[e*–*f] 1′ (cmp263) I*.
- DUP75 – *pgl-1(sam33[pgl-1::gfp::3xFLAG]) IV*.
- RAK146 – *sosi-1 eri-6[e–f] 1′ (cmp263) I; pgl-1(sam33[pgl-1::gfp::3xFLAG]) IV*.
- JMC231 **–** hrde-1(tor125[GFP::3xFLAG::HRDE-1]) III.
- RAK220 – *sosi-1 eri-6[e–f] 1′ (cmp263) I; hrde-1(tor125[GFP::3xFLAG::HRDE-1]) III*.
- AZ212 – *ruIs32 [pie-1p::GFP::H2B + unc-119(+)] III*.
- RAK218 – *sosi-1 eri-6[e–f] 1′ (cmp263) I; ruIs32 [pie-1p::GFP::H2B + unc-119(+)] III*.

### Male induction

L4s were plated on NGM plates, incubated at 30°C for 4, 5 and 6 hours and then placed at 20°C to recover. From the broods, males were isolated and mated with young adult hermaphrodites to maintain increased incidences of males within the populations.

### Brood size assay

Synchronized L1s of wild-type (N2) and *sosi-1 eri-6[e-f]Δ* worms were plated on NGM plates and cultured at 20°C. From those plates, individual L4 worms were isolated on NGM plates and cultured at 20°C (20 individuals per genotype). The number of eggs laid, and number of eggs hatched to produce progeny were counted for each hermaphrodite.

### Mating brood size assay

Synchronized L1s of *fog-2(oz40*) and male-containing populations of *sosi-1 eri-6[e-f]Δ* worms were plated on NGM plates and cultured at 20°C for a single generation. Mating plates were set up by pairing individual L4-stage *fog-2(oz40)* hermaphrodites with individual *sosi-1 eri-6*[e–f]*Δ* males grown at 20 °C (fifteen plates per cross). The number of progenies were counted for each hermaphrodite.

### Fluorescence microscopy

Synchronized L1s were plated and cultured on NGM plates at 20°C. L4-stage males were then selected and maintained for 24 hours at 20 °C without hermaphrodites to prevent mating. Whole animals were fixed in pre-chilled (–20°C) methanol for 5 minutes then washed twice in 1xPBST before being incubated in 0.17 mg/ml DAPI solution in PBST for 15 minutes, followed by three washes in 1xPBST. Imaging was performed on an AxioImager.M2 (Zeiss) using a Plan-Apochromat 63×/1.4-oil immersion objective. For sperm quantification, z-stacks were processed using FIJI^69,70^ and a projection of the stack capturing sperm in different focal planes were pseudocolored using the hyperstacks temporal-color code function. To count the number post-meiotic sperms in each male, the multipoint tool in FIJI was then used. The stack projection was pseudo-colored using Adobe Photoshop. Counts for post-meiotic sperm were taken for 15 biological replicates per genotype.

### Quantification of GFP fluorescence

Using the ‘measure’ function in FIJI^69,70^, the integrated density of 10 germline nuclei from the pachytene region of each worm was measured. Background signal was determined by taking the average of ten selected areas outside of the animals and the corrected total cell fluorescence (CTCF) of each germline nucleus was calculated as previously described^71^, with measurements obtained from 10 individual worms.

### RNA extraction

Synchronized L1s of wild-type and *sosi-1 eri-6[e-f]*Δ worms were plated on enriched peptone plates and cultured at 20°C. 8000 animals per sample were harvested as L4s (∼56 h at 20°C) for RNA extraction. Worms were washed off plates using water and then settled on ice to form a pellet. Water was aspirated off and worm pellets were resuspended in 1ml TRIzol reagent (Life Technologies) and freeze-thawed on dry ice followed by vortexing. Worm carcasses were pelleted using centrifugation and the supernatant containing RNA was collected. 0.2 volume chloroform was added to supernatant, mixed by vortexing, and the mixture was centrifuged. Then the aqueous phase was carefully transferred to a new tube, and samples were precipitated using isopropanol. Pellets were rehydrated in 50 μl nuclease-free H_2_O.

### mRNA-seq library preparation

Nuclease-free H_2_O was added to 7.5 μg of each total RNA sample, extracted from whole animals to a final volume of 100 μl. Samples were incubated at 65°C for 2 min then incubated on ice. The Dynabeads mRNA Purification Kit (ThermoFisher 61006) was used according to the manufacturer’s protocol. 20μl of Dynabeads were used for each sample. 100ng of each mRNA sample was used to prepare libraries with the NEBNext Ultra II Directional RNA Library Prep Kit for Illumina (NEB E7760S) according to the manual, using NEBNext multiplex oligos for Illumina (NEB E7335S). Library quality was assessed (Agilent BioAnalyzer Chip) and concentration was determined using the Qubit 1× dsDNA HS Assay kit (ThermoFisher Q33231). Libraries were sequenced on the Illumina NovaSeqX (SE 75-bp reads) platform. Three biological replicates were generated for wild-type (N2) and *sosi-1 eri-6[e-f]Δ* mutants cultured at 20°C.

### Small RNA library preparation

Small RNAs (18- to 30-nt) were size selected on denaturing 15% polyacrylamide gels (Bio-Rad 3450091) from total RNA samples. Libraries were prepared as previously described^72^. Small RNAs were treated with 5′ RNA polyphosphatase (Fisher NC0888901) and ligated to 3′ pre-adenylated adapter with Truncated T4 RNA ligase (Fisher 50-591-121). Small RNAs were then hybridized to the reverse transcription primer, ligated to the 5′ adapter with T4 RNA ligase (Fisher 50-811-609), and reverse transcribed with Superscript III (Thermo Fisher 18080–051) before being amplified using Q5 High-Fidelity DNA polymerase (Fisher NC0492039) and size selected on a 5% polyacrylamide gel (Bio-Rad 3450047). Library quality was assessed (Agilent BioAnalyzer Chip) and concentration was determined using the Qubit 1× dsDNA HS Assay kit (ThermoFisher Q33231). Libraries were sequenced on the Illumina NovaSeqX (SE 75-bp reads) platform. Three biological replicates were generated for wild-type (N2) and *sosi-1 eri-6[e-f]Δ* mutants cultured at 20°C.

### *In vitro* sperm activation assay

Synchronized L1s of wild-type and *sosi-1 eri-6[e-f]*Δ worms were plated on NGM plates and cultured at 20°C. L4 males from each strain were isolated on NGM plates without hermaphrodites and allowed to develop into adults at 20°C for 24 hours. 10–15 males were dissected in 15 μL sperm buffer (50mM HEPES, 50 mM NaCl, 25mM KCl, 5 mM CaCl2, 1 mM MgSO4, 0.1% BSA) with or without 400 ng/mL Pronase E (Millipore Sigma P8811) as described previously^73^. Sperm were released by gently nicking the tails, and the slides were incubated in a hybridization chamber for 15 minutes before a coverslip was mounted. Hermaphrodite sperm activation assay was done by isolating L4 stage hermaphrodites and allowing them to grow to young adults. For 30-40 hermaphrodite worms, sperm were released by making nicks close to the spermatheca. Sperm were imaged on an AxioImager.M2 (Zeiss) using a Plan-Apochromat 63×/1.4-oil immersion objective. Pseudopod formation was assessed for at least 200 sperm per genotype. Source Data is provided for pseudopod formation scoring.

### Bioinformatic analysis of mRNA-seq and small RNA-seq libraries

For small RNA libraries, sequences were parsed from adapters using Cutadapt v.4.1^74^ (options: -a TGGAATTCTCGGGTGCCAAGG -m 17 –nextseq-trim = 20 –max-n 2) and mapped to the *C. elegans* genome, WS258, using HISAT2 v.2.2.1^75^ (options: -q -k 11 -t -p 8) and the transcriptome using Salmonv.1.9.0^76^ (default options). Differential expression analysis was performed using DESeq2 v.1.38.0^77^.

For mRNA libraries, sequences were parsed from adapters using Cutadapt v.4.1^74^ (options: -a AGATCGGAAGAGCACACGTCTGAACTCCAGTCA -m 17 –nextseq-trim = 20 –max-n 2) and mapped to the C. elegans genome, WS258, using HISAT2 v.2.2.1^75^ (options: -q -k 11 -t -p 8) and the transcriptome using Salmonv.1.9.0^76^ (default options). Differential expression analysis was performed using DESeq2 v.1.38.0^77^. A log2(fold change) >1 or less than –1 and an adjusted P-value ≤0.05 was used as a cutoff to identify significantly up- and down-regulated genes in the DESeq2 output. Gene ontology enrichment analysis was performed using g:Profiler^78^.

Enrichment analyses used gene lists for RNAi pathways known to target sperm genes (CSR-1 target genes and PRG-1 target genes), previously identified histone genes and germline enriched genes (spermatogenesis-enriched genes, oogenesis-enriched genes, sex-neutral genes)^16,22,25,31,79-81^. Additional data analysis was done using R, Excel and Python. Venn diagrams were generated using InteractiVenn^82^.

