## Supplemental Figures and Tables for "Sensor-mediated fine-tuning of siRNA levels is required for spermatogenic piRNA pathway function"

#### SUPPLEMENTAL FIGURE 1

**A** **Enrichment analysis of genes up-regulated in *sosi-1 eri-6[e-f]*Δ L4 hermaphrodites**

| **Gene lists** | **$log_2$(enrichment)** | ***P*-value** |
|---|---|---|
| Spermatogenesis-enriched genes | 3.90 | $P = 4.3 \times 10^{-242}$ |
| Oogenesis-enriched genes | -INF | $P = 5.9 \times 10^{-10}$ |
| Sex-neutral germline genes | -0.54 | $P = 0.154$ |

**B** **Enrichment analysis of genes down-regulated in *sosi-1 eri-6[e-f]*Δ L4 hermaphrodites**

| **Gene lists** | **$log_2$(enrichment)** | ***P*-value** |
|---|---|---|
| Spermatogenesis-enriched genes | -0.44 | $P = 0.592$ |
| Oogenesis-enriched genes | 1.37 | $P = 1.9 \times 10^{-5}$ |
| Sex-neutral germline genes | 1.55 | $P = 3.5 \times 10^{-13}$ |

**Figure S1. (A)** Enrichment analysis for germline (sex-neutral, oogenic, and spermatogenic) genes amongst the genes up-regulated in *sosi-1 eri-6[e-f]*Δ L4s compared to wild-type animals is shown. -INF indicates there were no up-regulated genes overlapping with the gene list. **(B)** Enrichment analysis for germline (sex-neutral, oogenic, and spermatogenic) genes amongst the genes down-regulated in *sosi-1 eri-6[e-f]*Δ L4s compared to wild-type animals is shown. For **(A and B)**, two-tailed *P*-values for enrichment or depletion were calculated using the Fisher's exact test function in R.

### SUPPLEMENTAL FIGURE 2

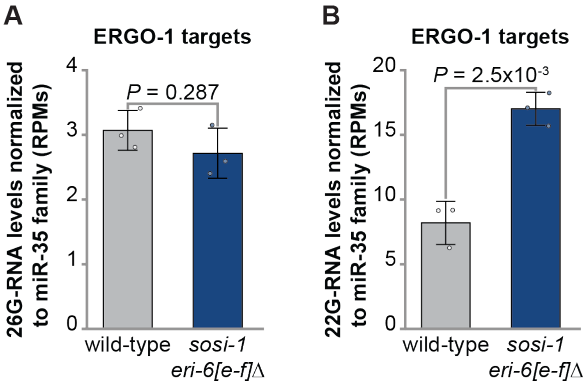


**Figure S2. (A)** 26G-RNAs and **(B)** 22G-RNAs mapping to ERGO-1 pathway target genes are counted, in reads per million (RPMs), for *sosi-1 eri-6[e-f]*Δ and wild-type L4 animals. Bar graphs represent the mean with dots representing summed RPMs for biological replicates and error bars indicating standard deviation. Statistical significance was calculated by two-tailed Welch's *t* test, mutants were compared to wild-type.

#### SUPPLEMENTAL FIGURE 3

**A** **Enrichment analysis of genes up-regulated in *sosi-1 eri-6[e-f]*Δ L4 hermaphrodites**

| Gene lists | $log_2$(enrichment) | *P*-value |
|---|---|---|
| PRG-1 targets | -0.54 | *P* = 0.049 |
| ALG-3/4 targets | 3.39 | *P* = $4.4 \times 10^{-229}$ |
| CSR-1 targets | -6.24 | *P* = $4.9 \times 10^{-33}$ |

**B** **Enrichment analysis of genes down-regulated in *sosi-1 eri-6[e-f]*Δ L4 hermaphrodites**

| Gene lists | $log_2$(enrichment) | *P*-value |
|---|---|---|
| PRG-1 targets | 0.80 | *P* = $3.6 \times 10^{-3}$ |
| ALG-3/4 targets | -0.93 | *P* = 0.091 |
| CSR-1 targets | 0.29 | *P* = 0.179 |

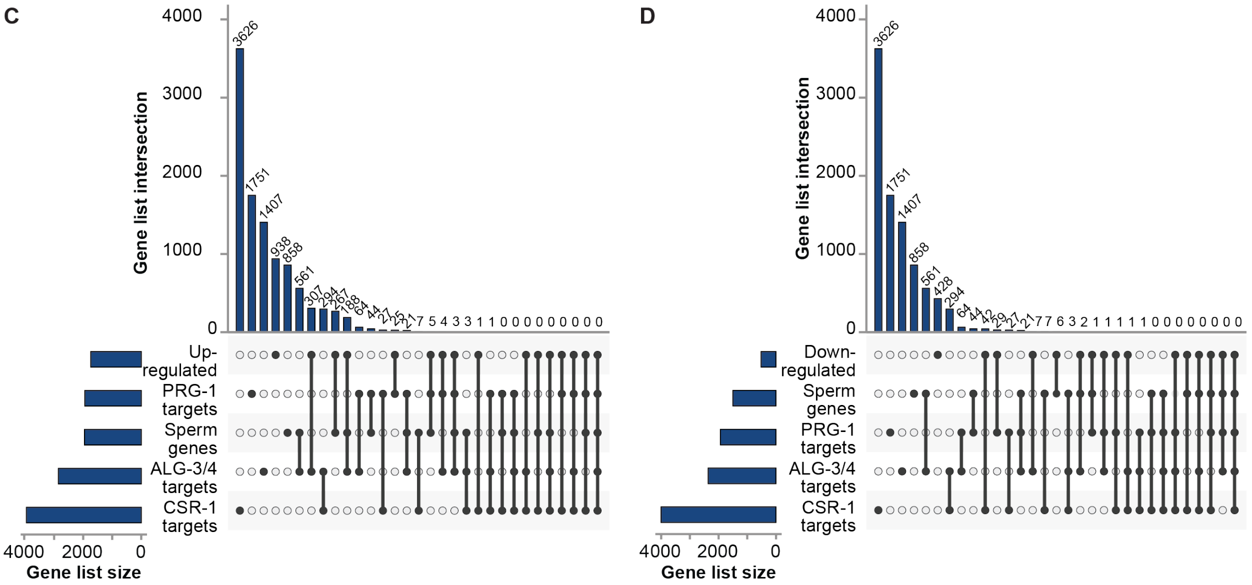


**Figure S3. (A)** Enrichment analysis for PRG-1, ALG-3/4, and CSR-1 pathway targets amongst the genes up-regulated in *sosi-1 eri-6[e-f]*Δ L4s compared to wild-type animals is shown. Two-tailed *P*-values for enrichment or depletion were calculated using the Fisher's exact test function in R. **(B)** Enrichment analysis for PRG-1, ALG-3/4, and CSR-1 pathway targets amongst the genes down-regulated in *sosi-1 eri-6[e-f]*Δ L4s compared to wild-type animals is shown. Two-tailed *P*-values for enrichment or depletion were calculated using the Fisher's exact test function in R. PRG-1, CSR-1, and ALG-3/4 pathways have overlapping target genes. Upset plots depicting the overlap of genes targeted by distinct small RNA pathways (ALG-3/4, CSR-1, and PRG-1 pathways), spermatogenic genes, and the genes **(C)** up- and **(D)** down-regulated in *sosi-1 eri-6[e-f]*Δ compared to wild-type L4 animals.

#### SUPPLEMENTAL FIGURE 4

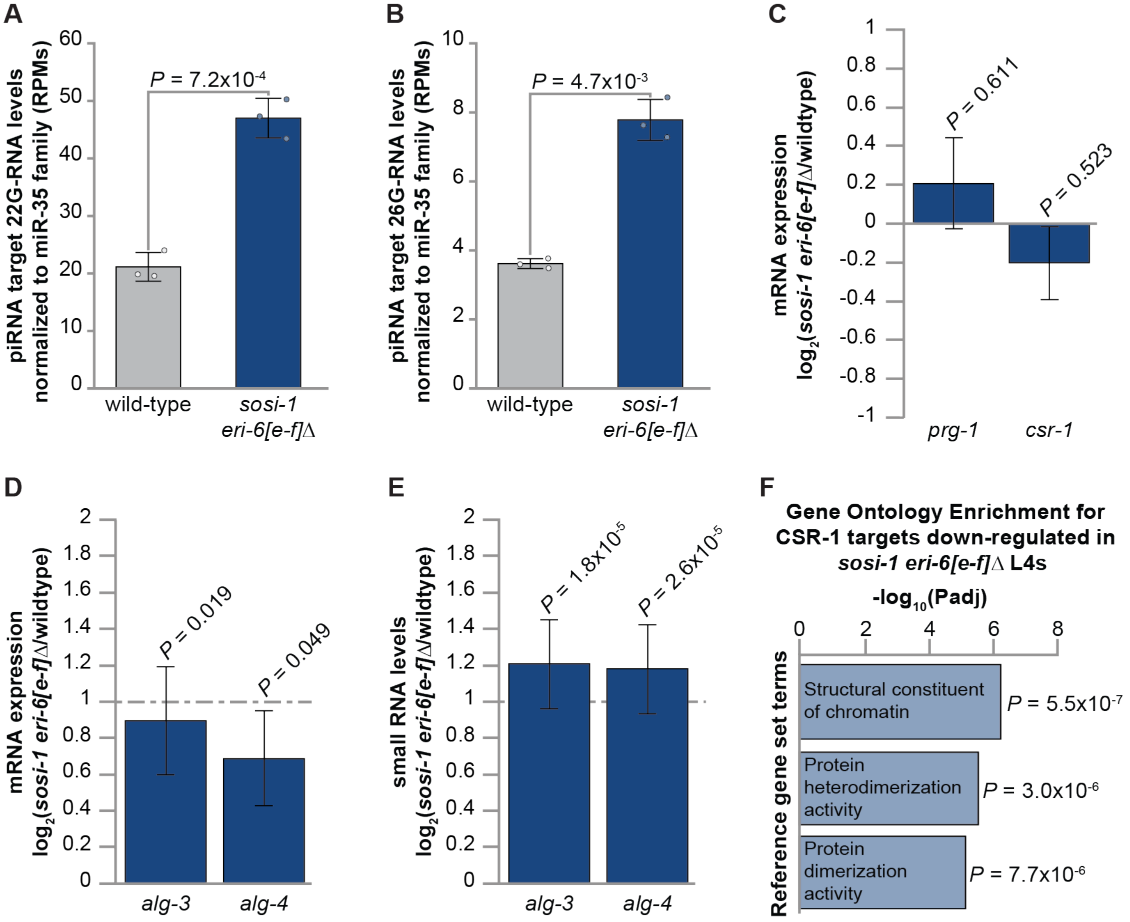


**Figure S4. (A)** 22G-RNA levels and **(B)** 26G-RNA levels normalized to *mir-35* family small RNA levels that map to all piRNA target genes are counted, in reads per million (RPMs), for wild-type and *sosi-1 eri-6[e-f]*Δ L4 hermaphrodites. Bar graphs represent the mean with dots representing summed RPMs for biological replicates and error bars indicating standard deviation. Two-tailed Welch's t-tests were performed to determine statistical significance. **(C)** Expression changes in *sosi-1 eri-6[e-f]*Δ L4s compared to wild-type animals for *prg-1* and *csr-1*. **(D)** Expression changes in *sosi-1 eri-6[e-f]*Δ L4s compared to wild-type animals for *alg-3* and *alg-4*. **(E)** Changes in small RNA levels in *sosi-1 eri-6[e-f]*Δ L4s compared to wild-type animals for all small RNAs mapping to *alg-3* and *alg-4*.

For **(C, D, and E)** Error bars indicate $log_2$(standard error) and *P*-value was calculated and corrected for multiple comparisons using the Benjamini and Hochberg method. **(F)** Shown is molecular function gene ontology enrichment for genes down-regulated in *sosi-1 eri-6[e-f]*Δ L4s that are CSR-1A, CSR-B, or CSR-1A+B targets.

#### SUPPLEMENTAL FIGURE 5

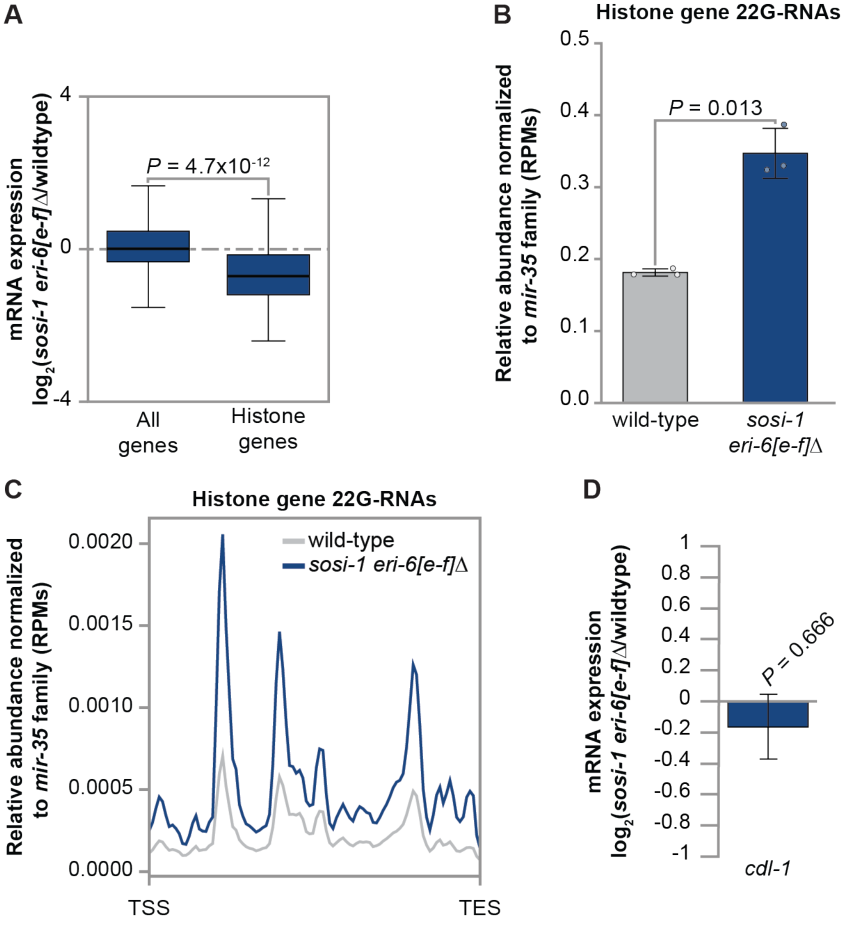


**Figure S5. (A)** Expression changes in *sosi-1 eri-6[e-f]*Δ L4s compared to wild-type animals for histone genes. Bolded midline indicates median value, box indicates the first and third quartiles, and whiskers represented the most extreme data points within 1.5 times the interquartile range, excluding outliers. Wilcoxon tests were performed to determine statistical significance between the $\log_2$(fold change) for histone genes compared to all genes, and *P*-value was adjusted for multiple comparisons. **(B)** 22G-RNA levels normalized to *mir-35* family small RNA levels that map to all hostone genes are counted, in reads per million (RPMs), for wild-type and *sosi-1 eri-6[e-f]*Δ L4 hermaphrodites. Bar graphs represent the mean with dots representing summed RPMs for biological replicates and error bars indicating standard deviation. Two-tailed Welch's t-tests were performed to determine statistical significance. **(C)** Metaprofile analysis showing the distribution of 22G-RNAs normalized to the *mir-35* family (RPMs) across histone genes in wild-type (gray) and *sosi-1 eri-6[e-f]Δ* (blue) L4 animals. The average of three biological replicates is shown. **(D)** Expression changes in *sosi-1 eri-6[e-f]*Δ L4s compared to wild-type animals for *cdl-1*. Error bars indicate $\log_2$(standard error) and *P*-value was calculated and corrected for multiple comparisons using the Benjamini and Hochberg method.

**SUPPLEMENTAL TABLES.**

**Supplemental Table 1.** Library mapping statistics.

| Library | Total Reads | Reads mapping to the WS258 genome | Reads mapping to WS258 mRNA, ncRNA, and pseudogenic transcripts | Number of 21U Reads mapping to the WS258 genome | Number of 22G Reads mapping to the WS258 genome | Number of 26G Reads mapping to the WS258 genome |
|---|---|---|---|---|---|---|
| HM_88_N2_PO_20C_L4_mRNA_1 | 38470201 | 37815896 | 24632746 | | | |
| HM_89_N2_PO_20C_L4_mRNA_2 | 34737871 | 34172613 | 20637972 | | | |
| HM_90_N2_PO_20C_L4_mRNA_3 | 41569109 | 40974875 | 27456019 | | | |
| HM_94_eri_6_e_f_sosi_1_PO_20C_L4_mRNA_1 | 31931083 | 31503328 | 23601520 | | | |
| HM_95_eri_6_e_f_sosi_1_PO_20C_L4_mRNA_2 | 61350203 | 60451462 | 46804222 | | | |
| HM_96_eri_6_e_f_sosi_1_PO_20C_L4_mRNA_3 | 40884882 | 40283743 | 29499374 | | | |
| HM_85_N2_PO_20C_L4_small_RNA_1 | 66204898 | 55814114 | 28080869 | 2653614 | 12699905 | 412155 |
| HM_86_N2_PO_20C_L4_small_RNA_2 | 63221960 | 55112460 | 20522225 | 3372319 | 9432673 | 457518 |
| HM_87_N2_PO_20C_L4_small_RNA_3 | 64765119 | 56080087 | 20550697 | 3574626 | 9451647 | 634512 |
| HM_91_eri_6_e_f_sosi_1_PO_20C_L4_small_RNA_1 | 68320186 | 59323435 | 23798095 | 4101379 | 11151216 | 834766 |
| HM_92_eri_6_e_f_sosi_1_PO_20C_L4_small_RNA_2 | 71336339 | 60988067 | 23077247 | 4250884 | 10431366 | 905975 |
| HM_93_eri_6_e_f_sosi_1_PO_20C_L4_small_RNA_3 | 66703269 | 56845415 | 22712346 | 3506740 | 10370826 | 710128 |
